# VARIATION AND SELECTION IN AXON NAVIGATION THROUGH MICROTUBULE-DEPENDENT STEPWISE GROWTH CONE ADVANCE

**DOI:** 10.1101/2020.01.29.925602

**Authors:** Stephen G Turney, Indra Chandrasekar, Mostafa Ahmed, Robert M Rioux, George M Whitesides, Paul C Bridgman

**Affiliations:** Department of Molecular and Cellular Biology, Harvard University, Cambridge, MA 02138; Department of Anatomy and Neurobiology, Washington University School of Medicine, St. Louis, MO 63110; Department of Chemical Engineering, Pennsylvania State University, University Park, PA 16802; Department of Chemistry, Pennsylvania State University, University Park, PA 16802; Department of Chemistry and Chemical Biology, Harvard University, Cambridge, MA 02138.

## Abstract

Myosin II (MII) activity is required for elongating mammalian sensory axons to change speed and direction in response to Nerve Growth Factor (NGF) and laminin-1 (LN). NGF signaling induces faster outgrowth on LN through regulation of actomyosin restraint of microtubule advance into the growth cone periphery. It remains unclear whether growth cone turning on LN works through the same mechanism and, if it does, how the mechanism produces directed advance. Using a novel method for substrate patterning, we tested how directed advance occurs on LN by creating a gap immediately in front of a growth cone advancing on a narrow LN path. The growth cone stopped until an actin-rich protrusion extended over the gap, adhered to LN, and became stabilized. Stepwise advance over the gap was triggered by microtubule +tip entry up to the adhesion site of the protrusion and was independent of traction force pulling. We found that the probability of microtubule entry is regulated at the level of the individual protrusion and is sensitive to the rate of microtubule polymerization and the rate of rearward actin flow as controlled by adhesion-cytoskeletal coupling and MII. We conclude that growth cone navigation is an iterative process of variation and selection. Growth cones extend leading edge actin-rich protrusions that adhere transiently (variation). Microtubule entry up to an adhesion site stabilizes a protrusion (selection) leading to engorgement, consolidation, protrusive activity distal to the adhesion site, and stepwise growth cone advance. The orientation of the protrusion determines the direction of advance.

## INTRODUCTION

Axon elongation has been characterized through modeling as a biased random walk that involves discrete growth steps (Katz et al., 1984). It has also been suggested that growth cones navigate using a spatial sensing mechanism (detecting a change in concentration gradient across the growth cone) as opposed to a temporal sensing mechanism (detecting changes in receptor occupancy over time) (Goodhill and Urbach, 1999; Mortimer et al., 2008). The view that growth cone turning is driven by an actin-based sensing and steering mechanism that involves stabilization of polarized protrusions as a first step (perhaps through actin bundling and adhesion) is widely accepted (Bentley and Toroian-Raymond, 1986; Davenport et al., 1993; Gomez and Spitzer, 1999; Kater and Rehder, 1995; Menon et al., 2015; Robles and Gomez, 2006; Zheng et al., 1996). Similar mechanisms have been proposed for neurite initiation (Dent et al., 2007), elongation (Suter and Miller, 2011), and for chemotaxis behavior by non-neuronal cells (Stephens et al., 2008). However, microtubule dynamics also correlate with growth cone movement (Sabry et al., 1991; Tanaka and Kirschner, 1991), initial neuronal polarization (Witte et al., 2008) and are required for turning (Buck and Zheng, 2002). Bulk advance of microtubules correlates with neurite elongation (Athamneh et al., 2017) and microtubule associated proteins have been implicated in steering (Pavez et al., 2019). In addition, it has been shown that microtubules or microtubule-based motors can be manipulated to influence turning even when actin dynamics are unperturbed (Challacombe et al., 1997) (Buck and Zheng, 2002; Kahn and Baas, 2016; Nadar et al., 2008). Thus, it remains unclear whether microtubule advance drives growth cone advance or vice versa.

To determine if actin cytoskeletal dynamics drives turning, we focused on the role of actin-dependent motor protein, myosin II (MII) in regulating growth cone direction and advance. Previous work showed that growth cone turning on laminin-1 (LN) at a border with poly-l-ornithine (PLO) is MII dependent (Turney and Bridgman, 2005) and also that MII may influence adhesion to LN (Ketschek et al., 2007). Furthermore, inhibition of MII prevents turning in response to inhibitory cues that do not induce collapse (Hur et al., 2011). In recent work we determined that NGF stimulates outgrowth through regulation of actomyosin restraint of microtubule advance (Turney et al., 2016). However, the common requirement for MII is also consistent with the possibility that growth cone preference for a substrate (i.e., turning) is partially a consequence of the relative degree of MII-dependent traction force pulling consistent with an actin-based steering mechanism as described by the molecular clutch hypothesis (Mitchison and Kirschner, 1988; Suter and Forscher, 2001). In addition it has been proposed that MII-dependent traction forces generated by the growth cone may stimulate elongation by axon stretching and intercalated growth (Suter and Miller, 2011). To distinguish between these possibilities, we devised a simple assay to force growth cones to advance in discrete steps. Neurons were grown in a microfluidic Campenot chamber with their axons exiting onto narrow lanes of LN that were widely spaced (to prevent crossing between lanes) and had non-adhesive gaps over which growth cones had to step in order to continue advance on LN, PLO or fibronectin (FN). We reasoned that if a growth cone could extend filopodia across a gap to form an adhesive contact on LN the slowing of retrograde flow and development of tension caused by adhesion-cytoskeletal coupling would induce stepwise growth cone advance through traction force pulling (Lamoureux et al., 1989). If crossing is blocked by inhibition of MII activity then we could infer that MII influences growth cone preference for LN primarily through traction force pulling supporting the prevailing view that substrate coupling is instructive (necessary and sufficient) for advance during directed growth and that the main function of MII is to develop pulling forces via the adhesion sites. However, if MII inhibition does not block advance then the core mechanism may be regulation of actomyosin restraint of microtubule advance as we recently proposed (Turney et al., 2016). Here we test between these possibilities.

## RESULTS

### Novel assay for analysis of stepwise growth cone advance

Mouse dorsal root ganglion (DRG) or superior cervical ganglion (SCG) neurons were cultured in a microfluidic Campenot chamber that served to hinder migration of glia into the axon compartment and to chemically isolate distal axon segments and growth cones from cell bodies. Axons exiting the channels of the chamber (10 μm wide) grew onto narrow LN lanes (9-12 µm wide). On many lanes we produced gaps (non-adhesive regions) over which axons had to cross for the elongation to continue. Both the lanes and the gaps were created by a new method of substrate patterning (Live Cell Substrate Patterning or LSCP; see Experimental Procedures) which uses region of interest (ROI) scanning of intense multiphoton laser light to remove PLO, FN and LN from the glass surface in well-defined patterns. The irradiated substrate regions could be distinguished from non-irradiated regions using reflected light laser scanning microscopy. The non-irradiated regions appeared darker in reflected light and supported adhesion and growth, while the lighter, irradiated regions did not (Figure 1).

**Figure 1.**
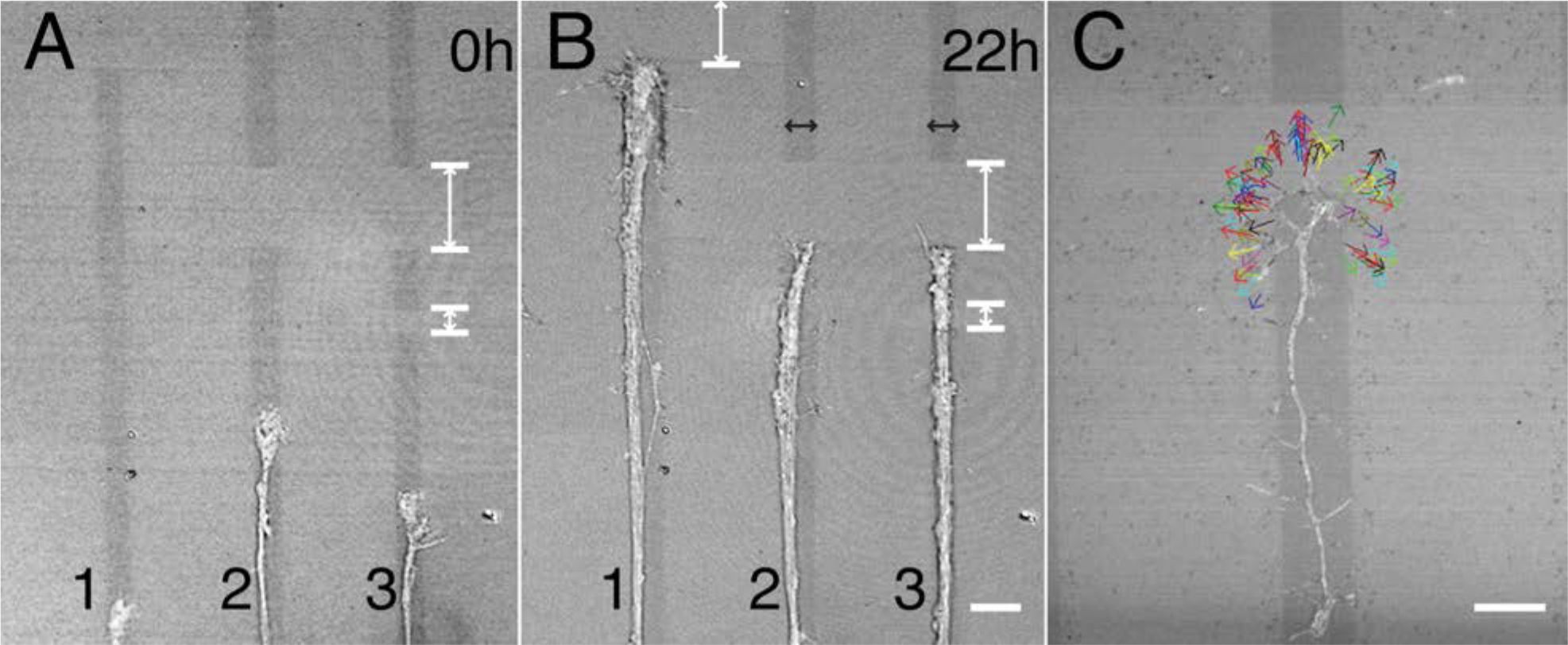
Axons elongating along LN lanes stop at non-adhesive gaps, but growth cone dynamics and mass addition continue. (A) Three axons growing on narrow LN lanes (dark grey stripes labeled 1-3) created by LCSP at the beginning of intermittent monitoring by DIC and reflected light imaging for 22 h. (B) All three axons have stopped elongating at the non-adhesive gaps (lighter areas within lanes indicated by double white arrow and bars) and remain stopped at 22 h. One axon (lane 3) crossed a short gap prior to stopping at the wider gap. Additional axons have grown along some of the lanes during monitoring (lane 1) but also stop at the gap. The axon calibers increase after 22 h (lanes 2, 3). Bar=18 µm (C) Growth cone protrusive activity continues when axons are stopped at gaps. Colored arrows show the filopodial extension positions from a time-lapse recording taken at 2 min intervals (see Movie S1). Bars=12 µm.

Reflected light was also used to determine close (adhesive-type) contacts of the protrusions with the substratum (Gomez et al., 1996). Gaps that were more than twice the lane width (∼10 µm) caused elongation to stop (Figures 1A and 1B). While stopped, growth cones continued to extend protrusions in multiple directions from the leading edge, but the protrusions appeared to have short lifetimes (see below) because frequently they did not adhere to the substrate. (Figure 1C; Movie S1). If elongation stopped for multiple hours, growth cones and proximal neurites appeared to enlarge presumably due to having increased mass. If the gap length was less than the lane width (<10 µm), growth cones could occasionally extend protrusions (i.e. filopodia) that were long enough to reach over the gap to contact the substrate on the other side. If a protrusion formed a contact on LN increasing its lifetime, the growth cone typically crossed after a short delay and continued to advance. For gap lengths ranging from approximately the lane width to twice the lane width, the timing of crossing appeared to vary stochastically. In a series of experiments individual axons exiting the channels of the chamber and entering the axon compartment were followed for up to a 22 h period as they grew along lanes (9-12 µm in width) and then stopped or stopped and then crossed gaps (Figure 1B and 2A). We monitored multiple axons in separate lanes by long-term time-lapse imaging to capture the rare gap crossing events. The cumulative number of gap crossings increased with time such that ∼80% of the total occurred within 16 h (Table 1).

**Table 1.**

|  |  |  |
| --- | --- | --- |
| Crossing frequency (% of total) (9-14 μm gap) (16 h)* | 81% | 26 |
| Crossing frequency (+Bleb) (16 h) | 87% | 23 |
| Rate of filopodial contact across gap (per h) | 0.9±0.3 | 10** |
| Rate of filopodia contact across gap (+Bleb) (per h) | 2.1±0.6 | 6** |
| Crossing failures (Ct) | 31% | 29 |
| Crossing failures (+Bleb) | 58% | 26 |
| Crossing frequency (+Nocodazole) (16 h) | 9.5% | 21 |
| Crossing frequency (+Bleb & Noc) (16 h) | 32% | 19 |
| Crossing failures (+Bleb & Noc) | 47% | 15 |
| Filopodia lengths Ct (uniform substrate; no gap) | 5.8±0.3# | 58 |
| Ct (at gap) | 7.5±0.4## | 89 |
| +Bleb (at gap) | 10.7±0.7### | 101 |
| +Noc (at gap) | 6.2±0.4#### | 94 |
| +Noc & Bleb (at gap) | 7.8±0.6##### | 142 |

Rates: Mean ± SEM.
\*Number of growth cones followed over time; first image taken at or approaching a gap, second image (of same growth cone) taken after approximately 16 h.
\*\*Number of growth cones analyzed, 3-4 h recordings each

### Stepwise growth cone advance requires adhesion and is facilitated by MII-dependent cytoskeletal coupling, but does not require traction force generation

To test whether traction force generation drives growth cone advance, we created LN lanes that terminated at a border with PLO or FN, often placing a gap between the two substrates (Figure 2). Growth cones advancing on LN stopped when they reached a border. The lanes were too narrow for the growth cones to turn or sidestep. At a border with PLO, they could remain stopped for at least 60 h (longest time point observed). Some protrusions spanning the gap were stable after making contact with PLO yet did not lead to further growth cone advance. At a border with FN, growth cone advance paused (typically 3-4 h) and then, after adhesive contact was made with FN, continued at a slower speed (Figure 2C, D). The above is consistent with the idea that the speed of growth cone advance is a function of the level of adhesion-cytoskeletal coupling on substrates that support formation of adhesion complexes. In previous work, we found that retrograde flow rates were lower on LN than on FN suggesting that coupling is stronger on LN (Turney et al., 2016). Retrograde flow rates were highest on PLO, and, accordingly, PLO does not appear to support adhesion-cytoskeletal coupling (Table 2). If traction force pulling is required for advance, then its contribution is likely to be larger on LN than on FN, and smallest on PLO (Turney et al., 2016).

**Figure 2.**
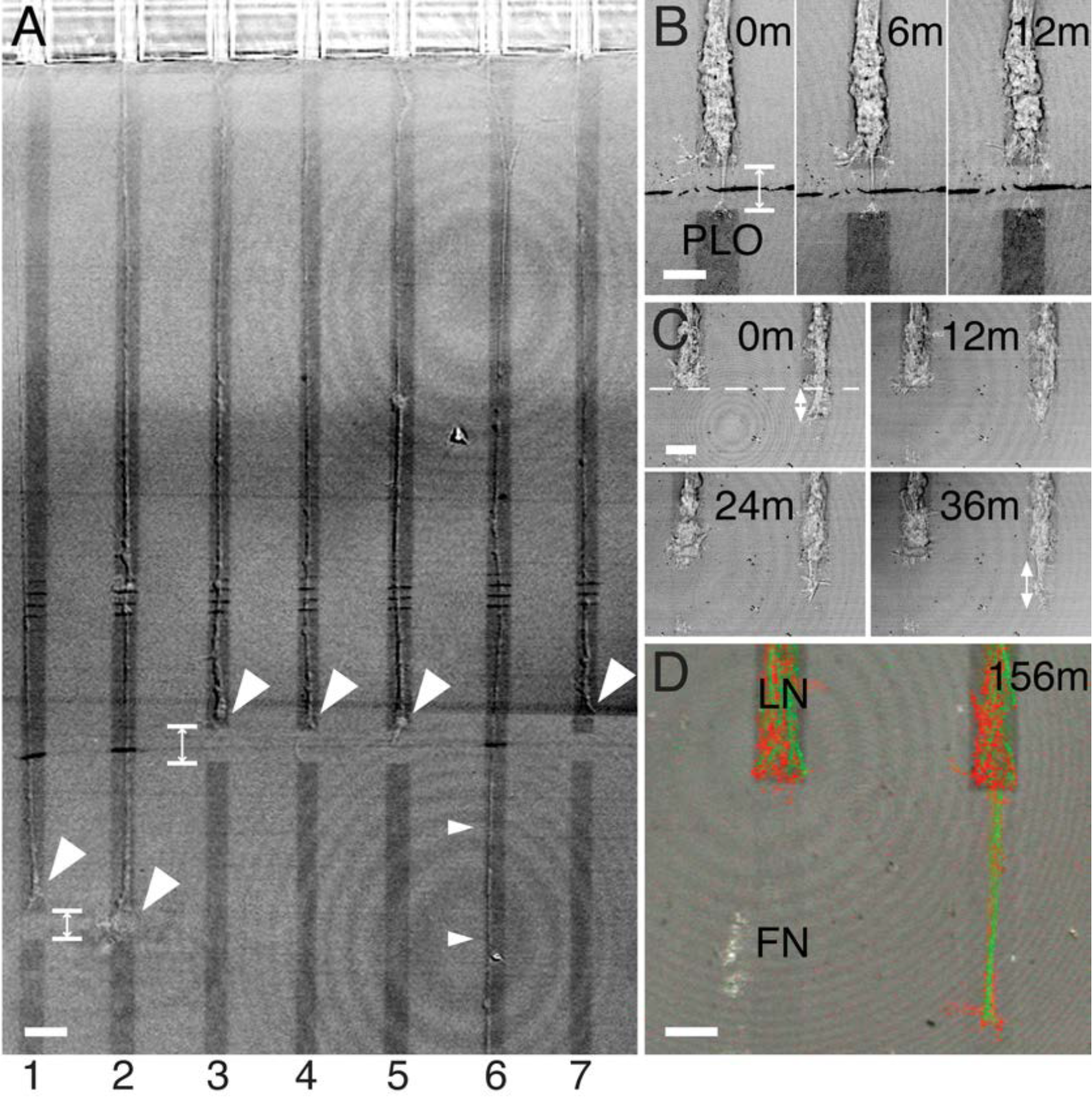
Combining patterning and microfluidics allows control of the local environment of elongating axons. (A) A low magnification view showing DRG axons growing out of the 10 µm-wide channels of a microfluidic Campenot chamber (top of field) into the axon compartment (aligned with lanes 1-7). The axon compartment substrate was patterned by LCSP to create LN lanes approximately 10 µm wide (dark grey stripes) separated by 40 µm-wide non-adhesive regions (lighter areas) as observed by merged reflected-light and DIC images. LCSP was used to create non-adhesive gaps on lanes 1-5 and 7 (double-ended arrows indicate the gaps on lanes 1 and 3). Axon elongation stopped at the 12 µm-long gaps (large arrowheads) but continued uninterrupted on lane 6 (small arrowheads). Bar=24 µm. (B) A sequence showing that growth cone advance on LN (lighter grey at top) toward PLO (darker grey at bottom) stops at a gap between the two apposed substrates. At 0 min, growth cone advance had been stopped for more than 1 h. The growth cone had extended a process and contacted PLO across the gap but did not advance. Bar=10 µm. (C) A sequence showing that growth cone advance on LN pauses at the interface (no gap) between LN and fibronectin (FN). At 0 min, advance had been paused for at least 2 h. The growth cone on the left remained stopped at the interface (dashed line), while the growth cone on the right slowly advanced. Bar=10 µm. (D) The same axons as in C following fixation and staining for actin with rhodamine phalloidin (red) and for microtubules with a Mab to tyrosinated tubulin (green) at the time indicated. Bar=10 µm.

**Table 2.** Parameter Rate (μm/min) N

|  |  |  |
| --- | --- | --- |
| Retrograde flow rate on<br>LN<sup>&</sup> | 3.1±0.3 | 64* |
| Retrograde flow rate on<br>PLO | 5.7±0.2<sup>#</sup> | 50** |

<sup>&</sup> Retrograde flow rates were analyzed as previously described (Turney et al., 2016)

Finally, we assessed crossing of gaps on LN lanes. Time-lapse observations revealed that growth cone advance occurred after a protrusion made adhesive contact with LN on the other side of the gap (Figure 3A). Formation of an adhesive contact usually led to further protrusive activity distal to the contact and then growth cone advance. The crossing events were rare (roughly 1 per h) primarily because most protrusions were too short (Table 1) and extended only part way over the gap (9-14 µm long). Thus, growth cones remained stopped for large periods of time until a sufficiently long protrusion formed to make contact with the substrate over the gap.

**Figure 3.**
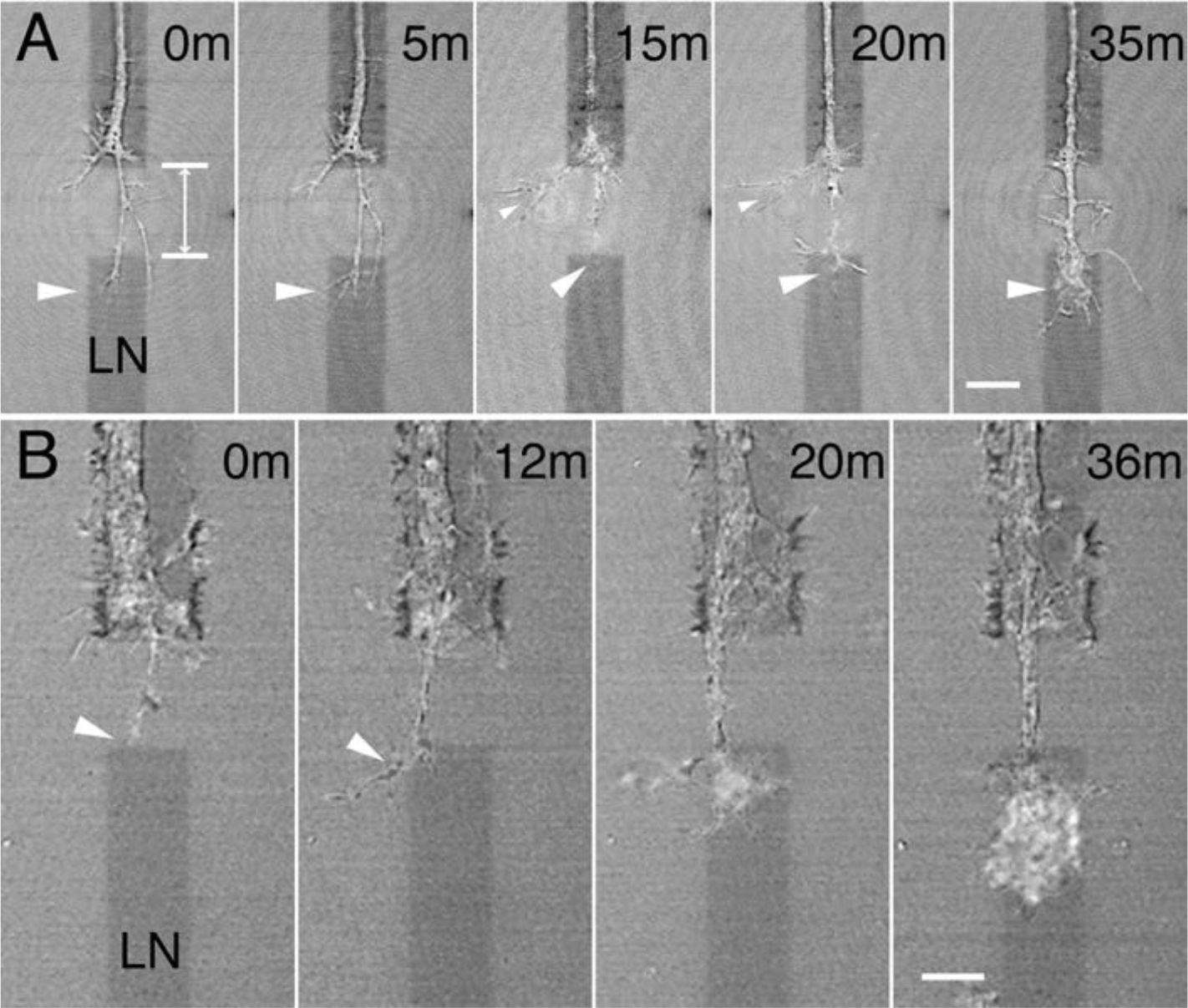
Control and blebbistatin-treated growth cones advance over non-adhesive gaps. (A) A sequence showing an untreated growth cone crossing a non-adhesive gap (double arrows). The growth cone extended filopodia to contact LN on the other side of a gap (arrowhead at 0 time). After contact a filopodium lengthened and branched (arrowhead at 5 min). At 15 min the filopodium making contact across the gap has partially retracted and no longer appeared to contact the post-gap LN (large arrowhead). Another filopodium has lengthened (small arrowhead). At 20 min contact with LN on the other side of the gap was re-established (large arrowhead). At 35 min the growth cone has crossed the gap (arrowhead). Bar=10 µm. (B) With blebbistatin treatment (50 µM in axon compartment only) growth cone behavior is similar to that of the control. A filopodium made initial contact with LN across the gap (arrowhead at 0 time). At 12 min the filopodium extended further (arrowhead at 12 min). At 20 min the expansion continued. By 36 min the growth cone has crossed the gap (see Movie S2). Imaging was by combined reflected-light and DIC at 800 nm. Bar=10 µm.

The lanes we created were straight and narrow, so growth cones could not turn but only advance by crossing a gap. Nevertheless, we suggest that growth cone behavior is fundamentally similar whether at a gap or at a naturally occurring decision point because it involves a pause in growth cone advance, exploration of the environment by protrusions, adhesion and then stabilization leading to further advance. Interestingly growth cone advance appears to pause in vivo at a decision point whether or not the growth cone crosses or it turns (Mason and Erskine, 2000). Thus, the same mechanism may underlie stepwise advance during decision point crossing and turning. One candidate is the actin-based steering mechanism involving MII dependent pulling force that aids in advance (Bridgman et al., 2001).

If a protrusion made contact with LN over a gap, it often persisted and became enlarged leading to growth cone advance. However, approximately 30% of the protrusions detached and retracted (Figure 3A; Table 1). The average lifetime of a protrusion was longer if it had adhered to LN than if it only extended over the irradiated substrate (i.e., had not adhered) (63% and 23% of adherent (N=22) and nonadherent (N=111) filopodia had lifetimes > 2 min, respectively). One possible explanation for the retraction is that the adhesion was not sufficiently strong to overcome MII dependent tension generated within the protrusion through a molecular clutch mechanism (Mitchison and Kirschner, 1988). Strong adhesion may be required for MII-dependent tension to pull the cytoplasm across the gap and allow elongation to continue. If tension is required to cross the gap then inhibiting MII may prevent advance. To test this possibility, we applied the MII inhibitor blebbistatin locally (in axon compartment only). Surprisingly the crossing of gaps was not eliminated. Blebbistatin (Bleb) is a specific inhibitor of myosin (mainly MII) and when used under appropriate conditions has direct effects only on myosin activity in mammalian cells (Allingham et al., 2005; Kolega, 2004; Limouze et al., 2004; Straight et al., 2003). For gaps of the same length, the frequency of crossing was roughly the same (81% vs 87% in 16 h; Table 1), indicating that MII dependent tension or traction force pulling is not required for crossing.

Time-lapse imaging revealed that growth cones crossing events at gaps were very similar with or without blebbistatin treatment (Figure 3B; Movie S2). However, blebbistatin treatment did cause a subpopulation of filopodia to increase in length (Figure 4A). These longer filopodia were more likely to make contacts with LN on the other side of a gap possibly aiding in crossing. Filopodia contacted LN on the other side of a gap approximately twice per hour in blebbistatin treated growth cones and once per hour in controls (Table 1). However, the number of crossing failures increased to close to 60% with blebbistatin treatment (Table 1) suggesting that the adhesion to LN or related processes were compromised leading to more failures. Therefore, the more frequent filopodial contacts were matched by an increased rate of failures resulting in a crossing frequency similar to that of controls.

**Figure 4.**
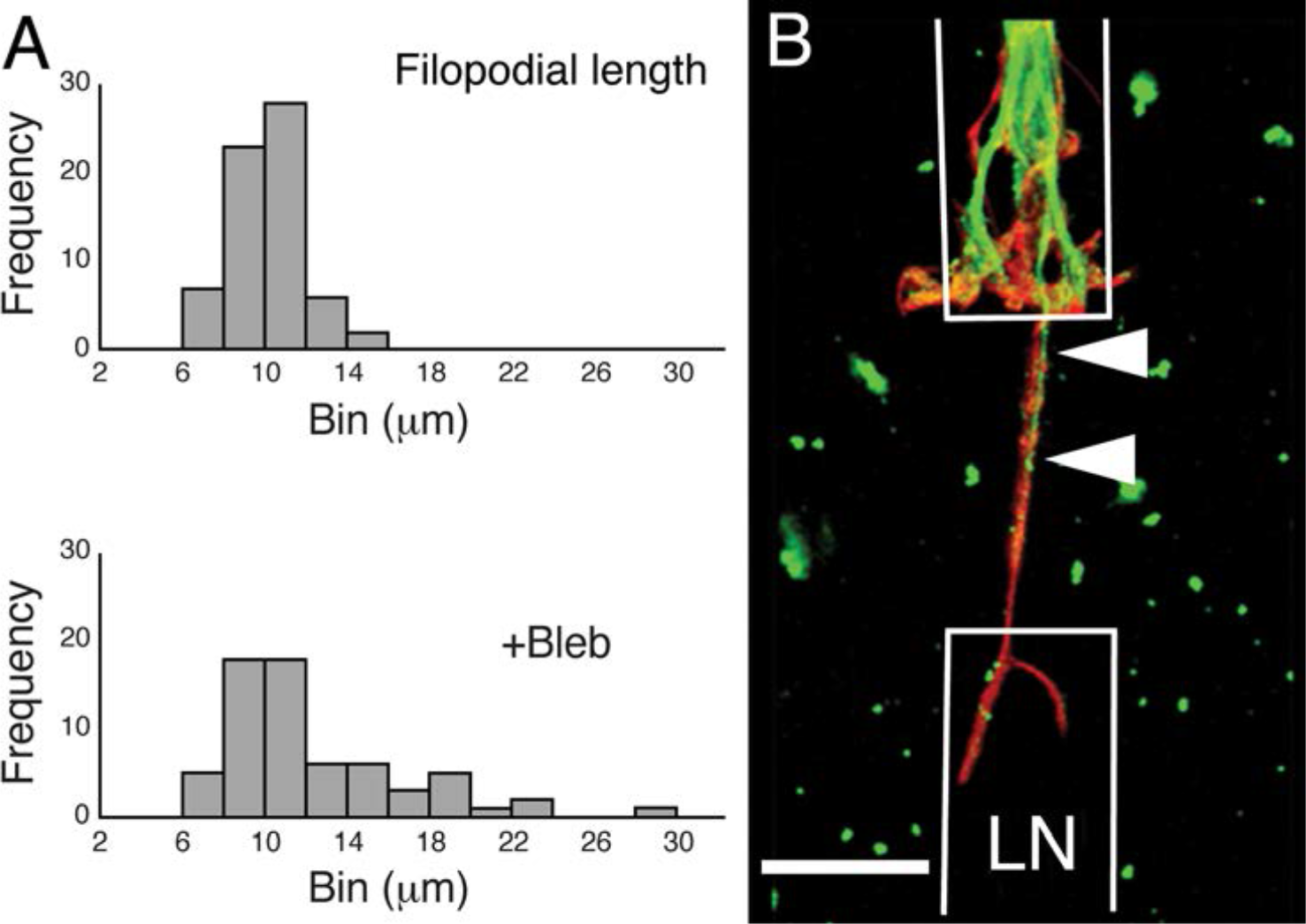
Filopodial characteristics of growth cones stopped at non-adhesive gaps. (A) Blebbistatin treatment (50 µM) induced the formation of longer filopodia. Top graph; the distribution of filopodia lengths measured from a 2 h recording (2 min intervals) of an untreated growth cone stopped at a non-adhesive gap. Bottom graph; the distribution of filopodia lengths recorded from a growth cone treated with blebbistatin. The distribution is skewed to the right indicating that more, longer-length, filopodia formed increasing the average length (Table 1). (B) An untreated growth cone stopped at a gap (white lines indicate the LN lane). Actin and dynamic microtubules were stained using rhodamine phalloidin (red) and a Mab to tyrosinated tubulin (green), respectively. Microtubules entered part way into the filopodium extending across the gap (arrowheads) Bar=12 µm.

### Growth cone advance correlates with microtubule entry into a protrusion

If advance over a gap does not require MII-dependent tension, then what drives advance when a protrusion makes adhesive contact over a gap? One possibility is that crossing only occurs if invading microtubules stabilize a protrusion. Consistent with this possibility, we stained for dynamic microtubules using an antibody to tyrosinated tubulin and found that microtubules were often present (>60%) in protrusions that had reached across gaps and formed adhesive contacts (Figure 4B). As a further test, we applied a low concentration of nocodazole (330 nM) to the axon compartment in order to interfere with microtubule polymerization. The concentration was calibrated to cause elongation to slow but not stop (Rochlin et al., 1996). Upon reaching a gap, growth cones remained active, grew in volume, but crossed only rarely during monitoring for up to a 36 h period (Figure 5A). Analysis of time-lapse images revealed that the failure to cross was mainly due to a decrease in the number of filopodia reaching over a gap and forming adhesive contact with LN. We found that the average length of filopodia was shorter in nocodazole (Noc) treated growth cones than in controls (Table 1). In 8.5 h of time-lapse recordings from five growth cones stopped at blocks, we captured only four instances of adhesive contacts. One of the four contacts led to crossing. The frequency of crossing determined from intermittent imaging (of a larger population) decreased to approximately 10% as measured over 16 h (Table 1). Washout of nocodazole led to increased filopodia length and a return to a higher frequency of crossing (Figure 6; 14 of 20 growth cones (70%) crossed by 16 h after washout). Low concentrations of taxol (100 nM) applied to the axon chamber had qualitatively the same effect as nocodazole at short times (<8 h) (Yvon et al., 1999), no crossing events were observed at 8 h (N=11). Although crossing was not observed at 16 h, we did not systematically study the effects of taxol further because growth cones sometimes slightly retracted or retreated from the gap with this longer treatment period. The higher crossing failure rate during nocodazole treatment suggests that dynamic microtubule advance may be required to stimulate extension of longer actin-rich protrusions perhaps because microtubules may enhance distal Rac1 activity by mediating the delivery or assembly of microtubule-bound Rac1 signaling complexes (Rochlin et al., 1999; Waterman-Storer et al., 1999).

**Figure 5.**
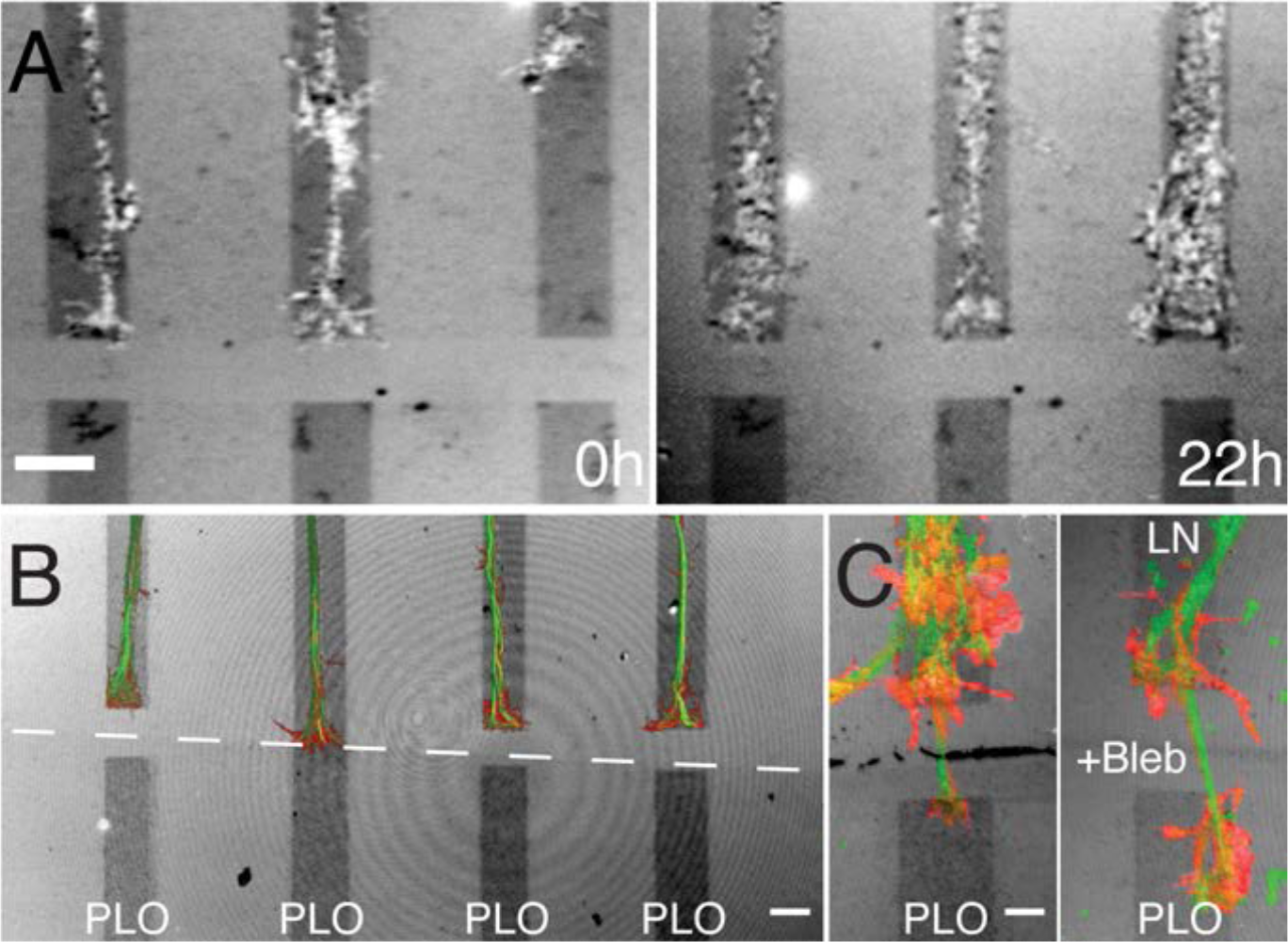
Nocodazole treatment suppresses growth cone crossing of non-adhesive gaps. (A) Axons on three adjacent LN lanes stopped at or approaching a non-adhesive gap (time 0h) immediately prior to adding nocodazole (330 nM). After 22 h (right panel) all three axons were still stopped at the non-adhesive gap. Bar=10 µm. (B) Untreated growth cones on LN stopped at borders with PLO (second lane from left) or with a non-adhesive gap immediately before PLO (remaining lanes). Axons were stained for actin (red) and dynamic microtubules (green). Axons reached the PLO border 5-6 h prior to fixation. Note that microtubules in the stopped growth cones rarely formed loops. Bar=10 µm. (C) Occasionally dynamic microtubules entered a process that reached across and made contact with PLO. However, the dynamic microtubules rarely extended into the portion of the filopodium on PLO (this is the same growth cone as in Figure 2B fixed 8 h after live imaging). Treatment with blebbistatin led to dynamic microtubules extending onto PLO and also increased expansion of actin-rich processes distally (right panel in C). Bar=5 µm.

**Figure 6.**
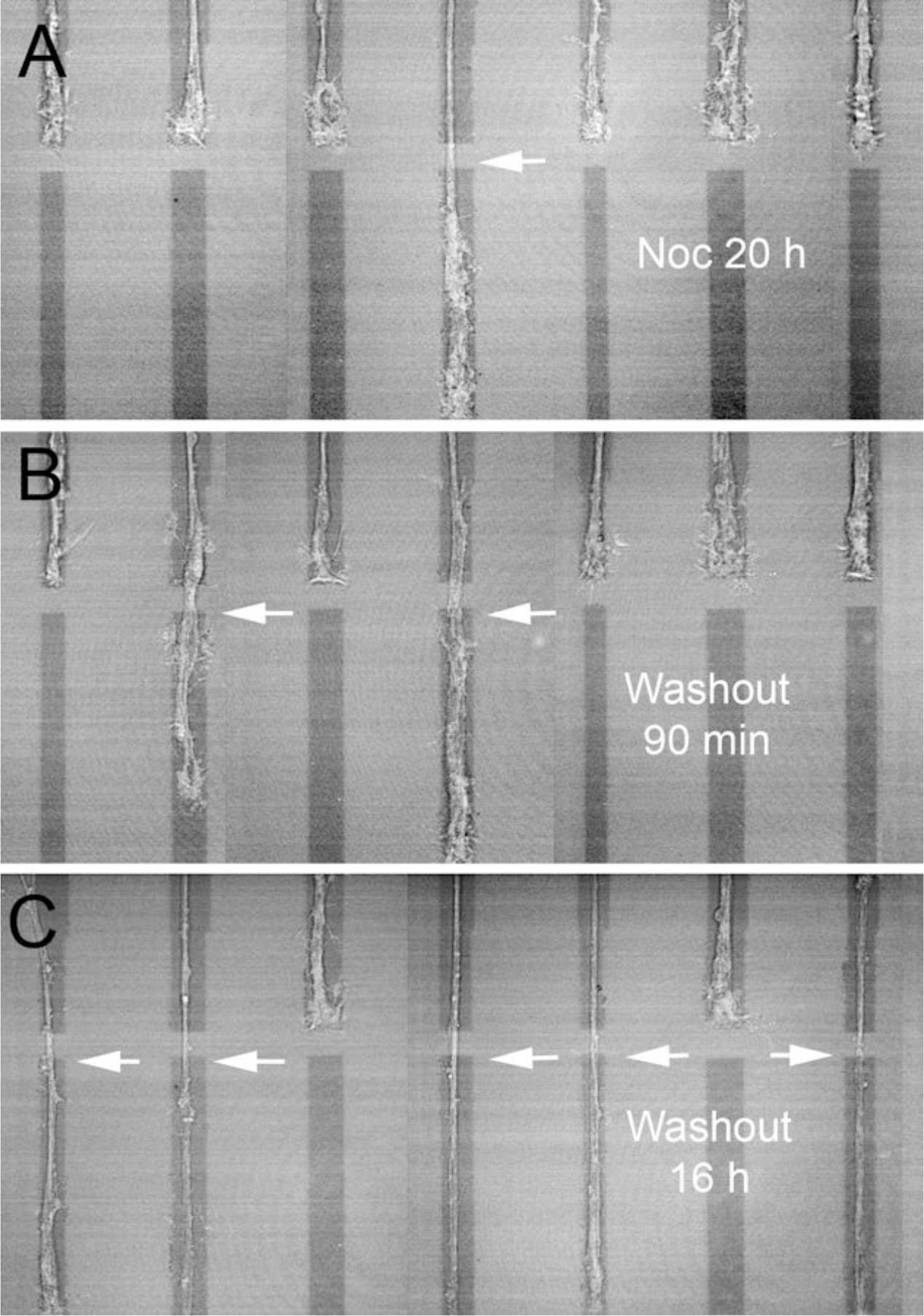
Washout of nocodazole leads to increased frequency of growth cone crossing of non-adhesive gaps. (A) Growth cones stopped at gaps after 20 h of nocodazole treatment. One crossing (arrow) has occurred. (B) Same growth cones 90 min after washout. One additional growth cone recently crossed a gap (left arrow). (C) Same area after 16 h after washout. A total of five growth cones have now crossed the gaps (arrows). (Note that stitching of images creates apparent vertical lines with varying positions in the background). Lanes are ∼ 10 µm in width.

To assess whether microtubule entry into protrusions was substrate and adhesion dependent, we fixed growth cones stopped at a border with the adhesive substrate PLO and stained them for dynamic microtubules and f-actin. Microtubules were only rarely (<5%) found in the portions of protrusions in contact with PLO (Figure 5B). On lanes that had a gap at the border with PLO, growth cones also stopped and over time produced protrusions that reached across the gap and contacted PLO. From time-lapse observations (30-60 min) these protrusions were rarely retracted indicating stable adhesive interactions, but they did not cause elongation to resume (see Figure 2B). After 1 to 8 h, we fixed and stained for actin and dynamic microtubules. A few of the protrusions (6 of 30 growth cones (20%)) contained microtubules extending either up to the PLO or only part way over the gap (Figure 5C, left panel). In contrast, when growth cones were treated with blebbistatin, the protrusions frequently (10 of 12 growth cones (>82%)) had microtubules extending over the gap and onto PLO (Figure 5C, right panel). In time-lapse recordings, F-actin containing protrusions were seen to make contact with PLO and grow in size shortly afterward. Over time, growth cones continued to advance on the PLO lanes. This is consistent with our previous finding that blebbistatin suppresses the ability of growth cones to alter their direction of growth at LN-PLO borders (Turney and Bridgman, 2005). These findings support the possibility that increased MII-dependent retrograde flow rates on PLO restrain dynamic microtubule entry thereby preventing advance.

### Retrograde flow rates regulate the probability of microtubule entry into a protrusion

MII driven retrograde actin flow has been shown to partially restrain the advance of dynamic microtubules into the actin rich periphery (Schaefer et al., 2002; Schaefer et al., 2008).

Inactivating MII decreases the restraint allowing dynamic microtubules to more readily invade actin-rich peripheral protrusions (Burnette et al., 2007; Turney et al., 2016). Thus, the reason blebbistatin treatment did not prevent crossing events, and actually increased their rate, could be that blebbistatin treatment enabled dynamic microtubules to more readily invade protrusions extending across gaps. Alternatively, blebbistatin may increase filopodial length independent of microtubule invasion. If the latter is true, then blebbistatin treatment may also facilitate crossing when microtubule dynamics are dampened with low concentrations of nocodazole. To test this possibility, we treated growth cones with both blebbistatin and nocodazole, applying these agents only to the axon compartment. We found that crossing events were partially restored (Figures 7A, B; Table 1). This finding is consistent with the idea that the longer filopodial lengths in response to blebbistatin treatment (Table 1) increases the probability of filopodial contact across the gap and even though microtubule polymerization rates are reduced the decreased restraint allows microtubules to invade (Turney and Bridgman, 2005) (Turney et al., 2016; Yang et al., 2012) (Ketschek et al., 2007).

**Figure 7.**
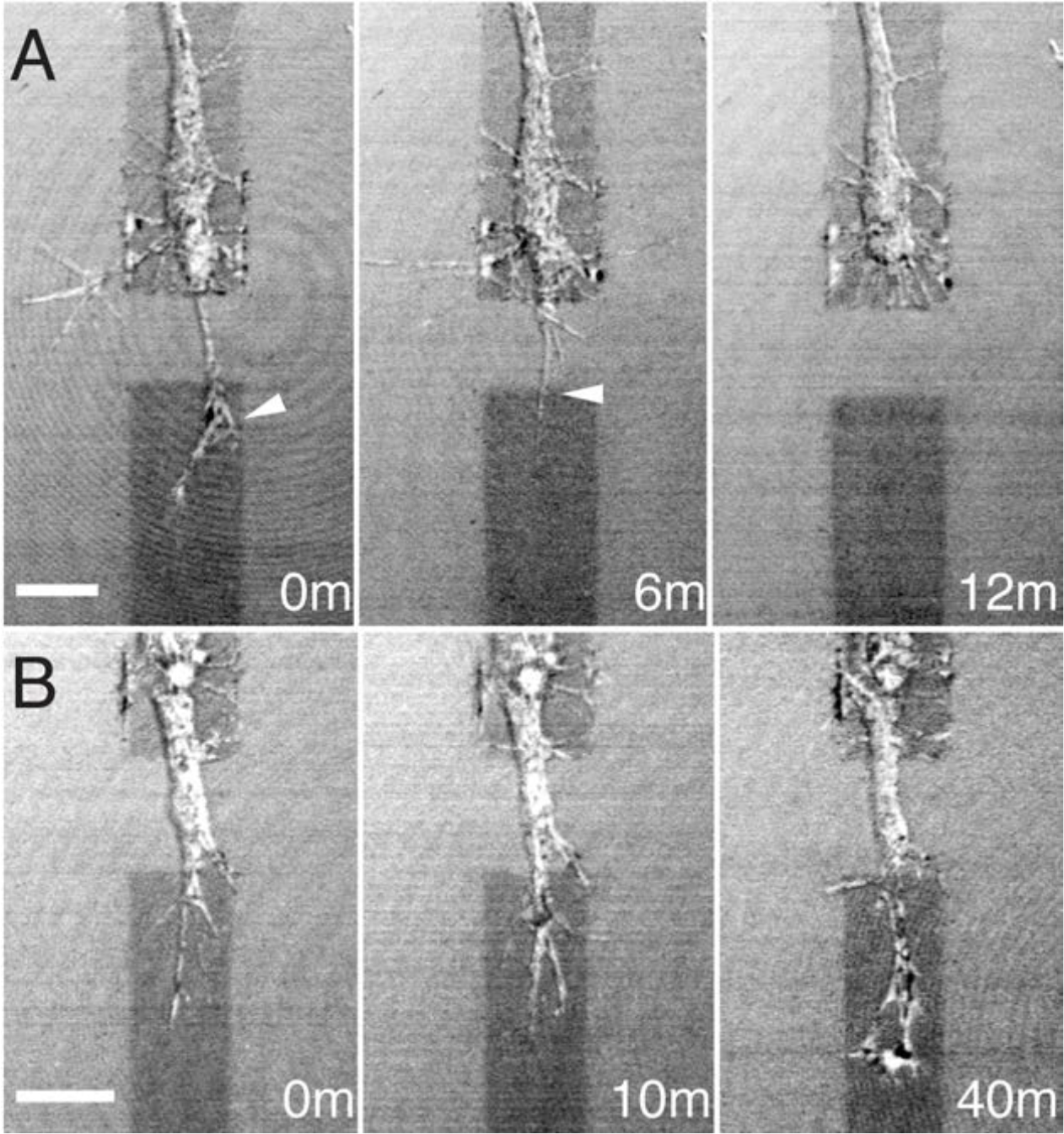
Treatment with a combination of nocodazole and blebbistatin partially restored growth cone crossing of non-adhesive gaps. (A) Treatment increased both the average filopodial length (Table 1) and the number of filopodial contacts across the gap (arrowheads). Bar=10 µm. (B) The frequency of crossing was lower than untreated growth cones, but was higher than for nocodazole treatment alone (Table 1). Images are combined reflected light and DIC at 800 nm. Time in minutes. Bar=10 µm.

To determine if retrograde flow rates differed in protrusions on LN versus PLO, we first measured the rates of retrograde flow in growth cones growing solely on one or the other substrate. Retrograde flow was measured by kymograph analysis of time-lapse image sequences from growth cones expressing GFP-LifeAct (or Ruby-LifeAct) (Fischer et al., 2006; Riedl et al., 2008; Turney et al., 2016). As mentioned above, retrograde flow in growth cones was significantly faster on PLO than on LN (Table 2). We then compared retrograde flow in growth cones stopped at LN-PLO borders. An abrupt transition in retrograde flow rates was detected between the filopodia in contact with PLO and the proximal regions of the growth cone on LN (Figure 8A). Normally retrograde flow in growth cones on LN or PLL slows gradually only in the transition zone (Medeiros et al., 2006; Turney et al., 2016; Van Goor et al., 2012; Yang et al., 2012). Thus, the retrograde flow rate can vary within individual growth cones depending on the underlying substrate. Contact with LN is more likely to induce adhesion complexes compared to contact with PLO (Nichol et al., 2016). When adhesion is coupled to the actin cytoskeleton via the molecular clutch, as occurs on LN, the retrograde flow rate decreases (Turney et al., 2016). The higher rate of retrograde flow in protrusions on PLO is likely to be more effective at restraining dynamic microtubules entry. Thus, axonal elongation is greatly decreased on PLO and stops at borders between the two substrates.

**Figure 8.**
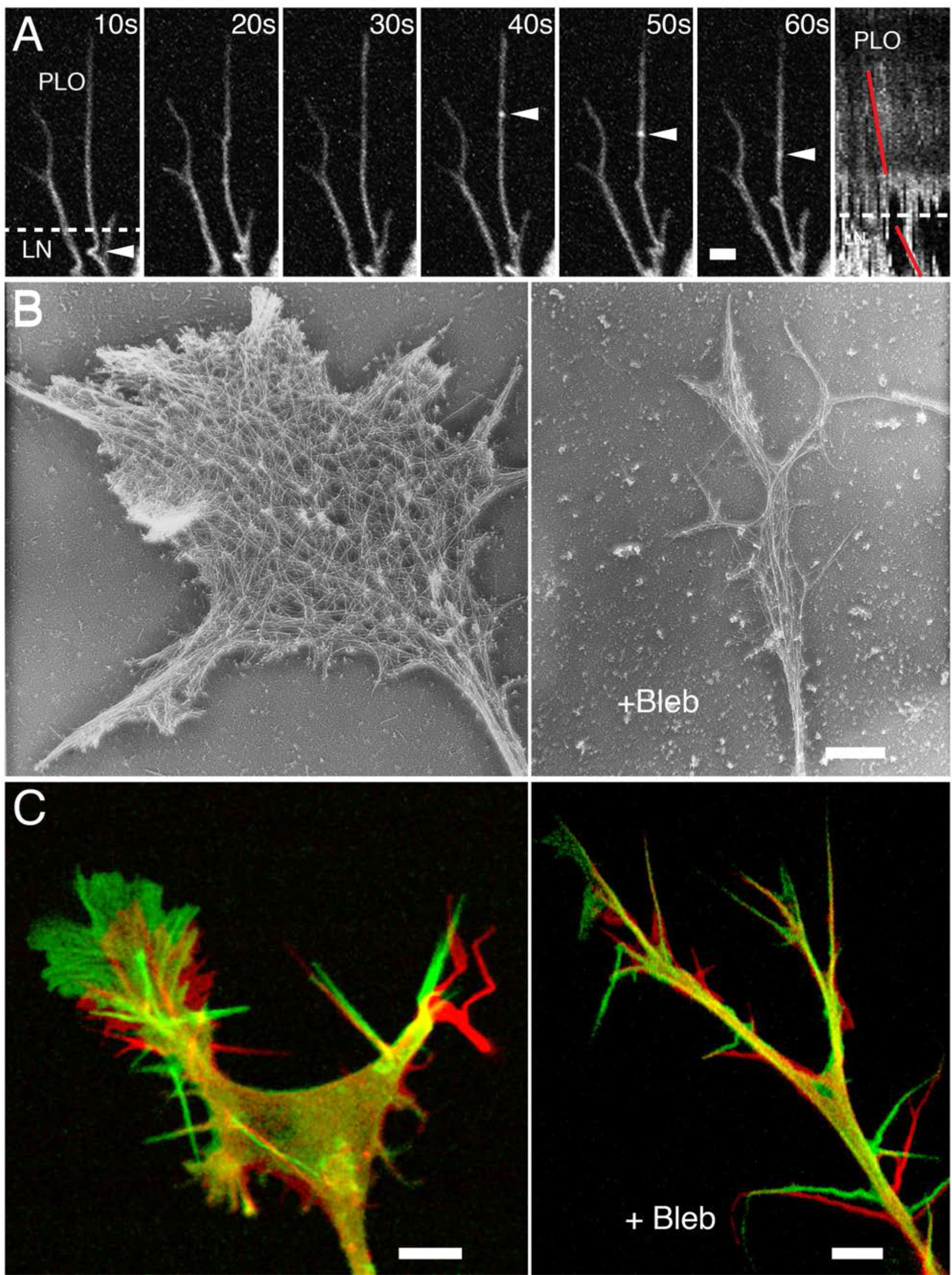
Changes to retrograde flow in individual protrusions adhering to LN and to PLO, and changes to actin organization and actin dynamics in growth cones after blebbistatin treatment. (A) A sequence showing retrograde flow of actin (GFP-LifeAct) at a LN-PLO border (time in seconds). Filopodia from a growth cone (below, out of the field) extend across the border. Bright actin particles (arrowheads at 40, 50 and 60 sec) move rearward with retrograde flow. Some buckling of a filopodium was seen to occur near the LN-PLO border at each time point (see arrowhead at 10 sec). Bar=2 µm. Rightmost panel: Kymograph generated from 40 successive frames of about the same region (shifted vertically). The different slopes (red lines) indicate faster and slower retrograde flow on PLO and LN, respectively. (B) Inhibition of MII activity with blebbistatin (Bleb) alters actin organization. In an untreated growth cone on LN (left panel), actin filaments are organized as bundles in protrusions at the leading edge (filopodia and lamellipodia) and in the proximal neurite. The large central domain has a dense meshwork of actin filaments. Following treatment with blebbistatin (> 1 h) the meshwork is lost and the central domain is greatly reduced in size (right panel). Actin filaments are arranged roughly in parallel all throughout the growth cone and neurite. (C) Actin dynamics in untreated (left panel) and blebbistatin-treated (right panel) growth cones on LN. Two successive frames (75 sec apart) were selected from the time lapse image sequence of each growth cone (see Movies S3 and S4) and displayed in red and green, respectively. In the untreated growth cone, actin-driven protrusion and retrograde flow were seen primarily at the leading edge of the branch on the left and aligned with the direction of advance. Following blebbistatin treatment (>45 min), growth cones were smaller in area and lacked a distinct central domain. Branching was frequent, and protrusion was no longer restricted to the leading edge of a single branch. Short-lived lamellipodia were extended from the periphery of both branches. Filopodia persisted and were also extended from the trailing neurite. (Note the shift in position of the branch point indicates forward advance of the neurite shaft.) Retrograde flow appeared to be more global and was associated with both branches and the neurite. Bars=6 µm.

Blebbistatin treatment has been shown to slow retrograde flow in Aplysia growth cones on poly-l-lysine (PLL) (Medeiros et al., 2006) and LN (Yang et al., 2012). To determine its effect in mammalian growth cones on LN, we used rotary shadowing electron microscopy (EM) and time-lapse imaging of GFP- or Ruby-LifeAct fluorescence to assess actin organization and retrograde flow, respectively. Similar to the Medeiros et al. results for PLL and Yang et. al. results for LN, blebbistatin treatment did not cause retrograde flow to stop presumably because actin treadmilling continues to drive the flow; however, after 30 min, the lamellipodia and the central domain became thin and finger-like. The actin meshwork of the central domain was largely eliminated (Figure 8B). Protrusive activity was no longer restricted to the leading edge as is typically observed in untreated controls undergoing expansion (Figure 8C). The character of retrograde flow changed from being distinct primarily in the periphery to being distinct in all portions of the growth cone and in the proximal neurite (Movies S3 and S4). The rate was difficult to measure accurately using kymographs because the direction of flow fluctuated wildly immediately after treatment and the growth cone morphology underwent rapid ongoing changes thereafter. The character and rate of retrograde flow after blebbistatin treatment appeared to be qualitatively the same on PLO as on LN. As has been previously observed, growth cone polarity and consolidation of the neurite were abnormal (Loudon et al., 2006). Filopodia persisted and transient lamellipodia formed along neurite branches and filopodia. Blebbistatin also alters adhesion complexes, actin organization and bundling on LN (Burnette et al., 2011; Goeckeler et al., 2008; Turney et al., 2016). The overall effect may be to reduce differences in bundling and retrograde flow on the two substrates thereby roughly equalizing the restraint of microtubule advance. As previously observed, axon elongation increased on PLO following blebbistatin treatment (Ketschek et al., 2007; Turney and Bridgman, 2005). The faster advance was presumably due to decreased restraint. On narrow LN lanes that terminated at a border with PLO (sometimes with a non-adhesive gap at the border), axon elongation stopped at the border and resumed on PLO only after blebbistatin treatment (9/9). Growth cones continued their advance on PLO even if it required turning in response to a non-adhesive border (Figure 9).

**Figure 9.**
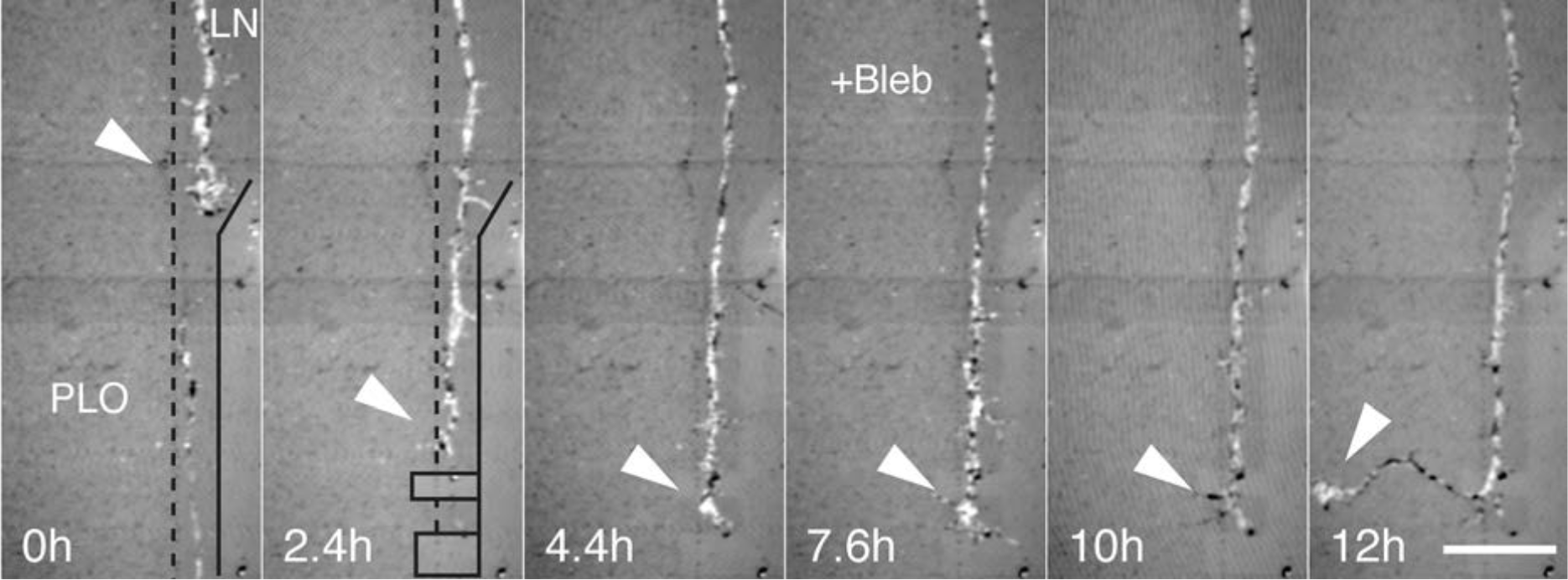
Growth cone advance on a LN lane bounded by PLO and a non-adhesive region was stopped by a non-adhesive gap, but turned and resumed advance on PLO after inactivation of MII. At 0 h, a mouse SCG neurite extended at 20 μm/h on a 7.5 μm-wide path of LN bounded by PLO (dashed line) and by a non-adhesive region created using LCSP (solid black line). At 2.4 h, the LN substrate was irradiated immediately in front of the growth cone to create barriers to advance (black boxes). The irradiation also ablated a second neurite growing in the opposite direction. By 4.4 h, the growth cone crossed the first (4 μm) gap and reached the second (8 μm) gap where its advance was blocked. The growth cone remained on the 7 μm-long region of LN between the two gaps rather than turning and advancing on the growth permissive PLO substrate. At 7.6 h, the MII specific inhibitor, blebbistatin, is added to the growth medium. By 10 h, the growth cone has turned and has advanced onto PLO. At 12 h, neurite outgrowth continued at approximately 12 μm/h on PLO. Imaging was by 800nm reflected light only. Bar=20 µm.

### Decreased probability of microtubule entry into protrusions leads to aversive turning and stopping of the growth cones

To assess whether the stopping of growth cone advance on LN at a border with PLO correlates with a low probability of microtubule entry into protrusions on PLO, we imaged dynamic microtubules using the (+) plus-end tracking protein GFP-EB3 (Figure 10; Movie S5) (Stepanova et al., 2003). GFP-EB3 was observed to penetrate occasionally into regions of broad lamellipodia on PLO but not in filopodia on PLO. The microtubule entry was not seen to produce enlargement of either the lamellipodia or the filipodia consistent with growth cone advance being fully stopped. The lifetime of the GFP-EB3 spots in lamellipodia was short as measured from time lapse recordings of four growth cones (22±3 s) and approximately the same as that observed in growth cones advancing slowly on LN and FN in low NGF (Turney et al., 2016). From the above, we conclude that a growth cone advancing on a narrow LN lane stops at a border with PLO in part because of the low probability of microtubule entry into protrusions on PLO. Other contributing factors may include decreased dynamic microtubule lifetimes and the interplay between retrograde flow and actin polymerization.

**Figure 10.**
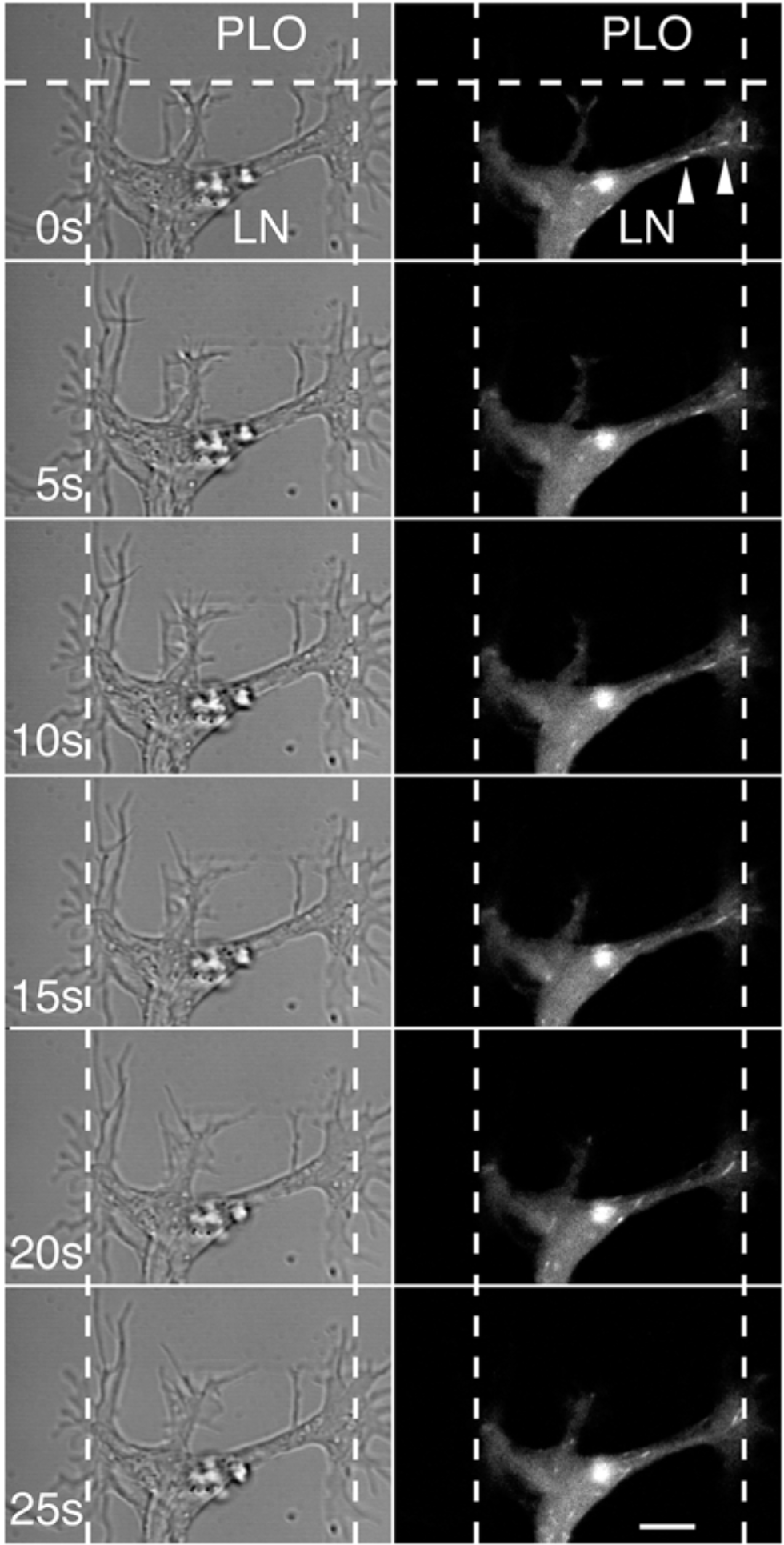
GFP-EB3 dynamics at a border between LN and PLO. A series of images (at 5 s intervals) from a time lapse recording (DIC and fluorescence; see Movie S5) showing the dynamics of EB3 relative to the lane boundaries (dashed vertical lines) and the border with PLO (dashed horizontal line). EB3 did not enter processes that crossed the border to contact PLO and rarely extended into portions contacting the non-adhesive lane boundaries. Bar=4 µm.

Growth cone turning at LN-PLO borders may be a consequence of the probability of microtubule entry into protrusions being lower on PLO than on LN. The difference in probability may be related to the difference in retrograde flow rates; however, it also possible that LN may stimulate or stabilize protrusions thereby increasing the probability of microtubule entry. To test this possibility, we compared the total numbers of filopodia forming on each substrate during turning at a border between LN and PLO. More filopodia formed on LN than PLO, but had longer detectable lifetimes on PLO primarily because the lamellipodia advance that was observed to engulf filopodia on LN was absent on PLO (Figure 11A and E). This resulted in persistent advance along borders between LN and PLO. In the same growth cone, we also compared the total numbers of filopodia during turning at a border between LN and a non-adhesive region produced by LCSP laser irradiation. Filopodia contacts to non-adhesive regions were greatly reduced compared to LN (Figure 11B and D). Growth cones on LN that turned at borders with non-adhesion regions advanced and grew away from the border shortly after turning. Inactivation of MII by blebbistatin treatment did not eliminate turning at non-adhesive regions (as it does for PLO), but after turning advance continued along the borders for long periods of time (Figure 11C). Therefore, turning is not likely to result just from enhanced filopodial formation induced by LN or from longer filopodial lifetimes. Instead, the increased probability of microtubule entry into protrusions, and stability that leads to further protrusion on LN is likely to be due to differences in the degree of restraint.

**Figure 11.**
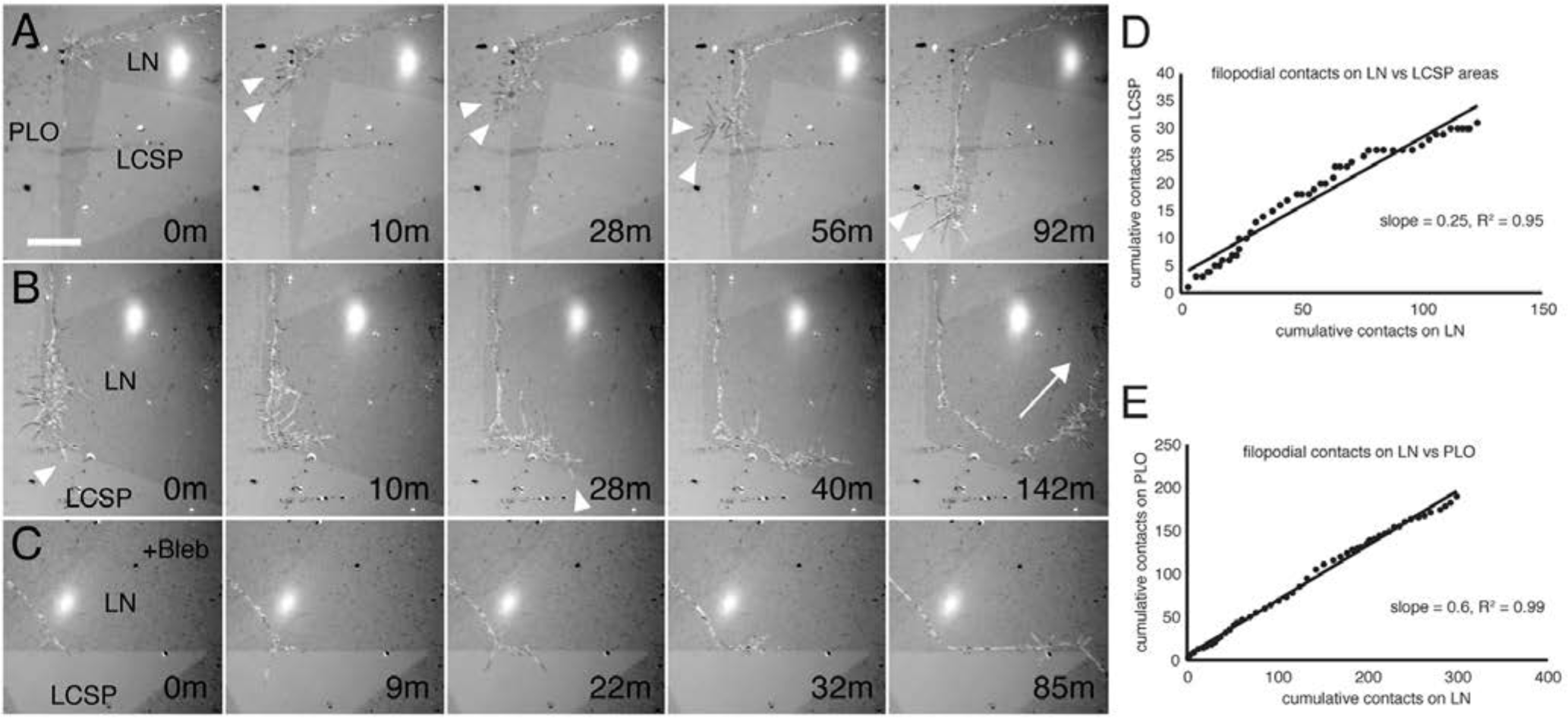
Filopodial dynamics during growth cone turning and advance on LN at borders with PLO and non-adhesive areas created by LCSP. (A) A growth cone advancing on LN approached a border with PLO. The growth cone turned to remain on LN even though it extended filopodia that made adhesive contact with PLO (dark filopodia indicated by arrowheads in the reflected-light images). As the growth cone continued along the LN-PLO border, it made frequent filopodial contact with PLO but did not cross onto PLO. (B) The growth cone reached a border with a LCSP area and turned to remain on LN. The filopodia extended over the LCSP area were transient and non-adhering (light filopodia indicated by arrowheads). The growth cone continued on LN but turned away from the LCSP border starting at 40 min (arrow at 142 min). (C) After blebbistatin treatment to inactivate MII, a growth cone on LN turned and advanced along a border with a LCSP area. (D) Number of filopodia extended during growth along a LN-LCSP border (border at top, prior to sequence shown in A). Fewer filopodia contacts were seen on the LCSP area than on LN as determined by linear regression analysis (slope=0.25). (E) When the same growth cone reached a border with PLO (sequence shown in A), it turned and grew along the border, consistently making more contacts on LN than on PLO (slope=0.6). Contacts to PLO were detectable for longer times. The linear relationships in D and E are representative of four additional recordings (not shown). Imaging was by 800 nm reflected-light optics only. Bar=18 µm.

To determine if dynamic microtubule entry into protrusions extending across non-adhesive gaps is necessary for crossing, we also monitored GFP-EB3 dynamics in growth cones that were stopped at the gaps. GFP-EB3 was observed to penetrate into protrusions that had reached across gaps to contact LN (Figure 12; Movie S6). Although the number of observations is small (due to the low probability of “catching” the rare crossing attempts in growth cones with bright fluorescence), notably there was a correlation between the depth of penetration and the chance of successful crossing event. If GFP-EB3 penetrated sufficiently far that it likely interacted with the putative adhesion site that had formed on the post-gap LN, then crossing ensued (four examples). However, if GFP-EB3 penetrated only part way into a protrusion (i.e., only into the portion overlying the gap) then crossing failed and the protrusion was retracted (three examples). This is consistent with our much larger number of time-lapse observations on crossing failures (Table 1). This result suggests that penetration of dynamic microtubule ends well into a protrusion is needed for stabilization on LN and this is likely to be necessary for advance of the growth cone through continued protrusion.

**Figure 12.**
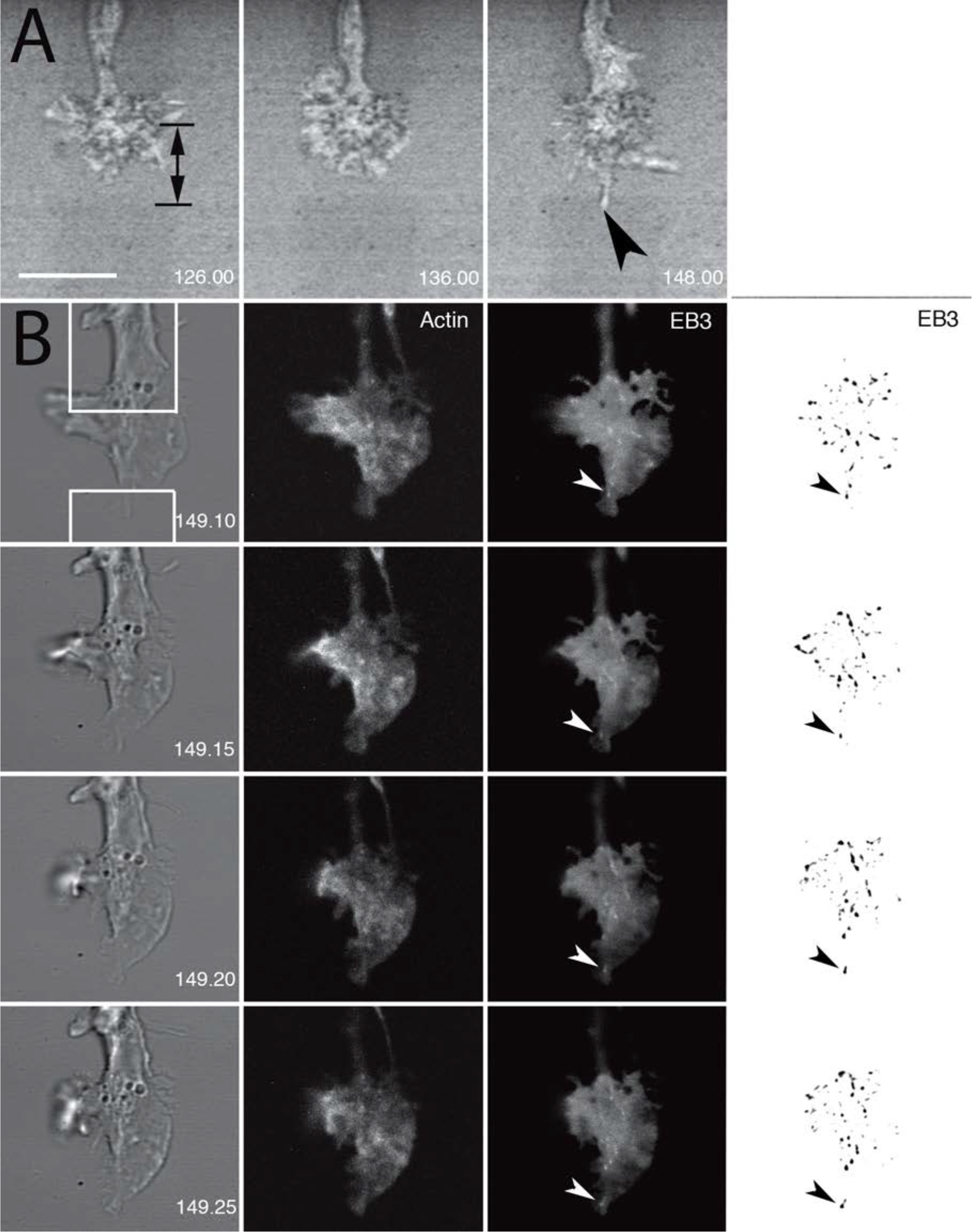
Growth cone advance requires microtubule invasion of an adherent filopodial protrusion. (A) A growth cone stopped at a non-adhesive gap (black double-ended arrow) on a narrow LN path was imaged using time lapse microscopy (simultaneous reflected-light and DIC optics, 2 min interval) until filopodial contact (arrowhead) was made with LN across the gap (148 min). Bar=12 µm. (B) The field was shifted and the scan zoom was increased by 10%. The growth cone was then imaged every 5 s using simultaneous DIC (left panels) and epifluorescence optics (middle panels: Ruby-LA, right panels: GFP-EB3). A dynamic microtubule invaded deeply into the adherent protrusion (white arrowheads; see Movie S6). Panels on far right show the EB3 spots in reversed grayscale (black arrowheads) after using a filter (SpotTracker plugin, ImageJ) and thresholding to enhance the punctate signal. Arrowhead indicates the same EB3 spot as in the panels immediately adjacent. The growth cone proceeded to advance over the gap. In other recordings, dynamic microtubule (+) ends that terminated advance before reaching the site of contact with LN (invaded only part way) failed to trigger crossing.

## DISCUSSION

The results of the current study are consistent with growth cone advance occurring in a stepwise manner. We found that growth cones advancing on narrow LN paths always stopped at a border with PLO, which is adhesive but does not support cytoskeletal coupling. To test whether the stopping is due to a balance of traction force pulling that is greater on LN than on PLO (or FN), we created non-adhesive gaps over which a growth cone could extend a filopodial protrusion. In the absence of MII activity (that is, with blebbistatin treatment), crossing occurred onto all three substrates tested (LN, FN, PLO). Crossing occurred only after a protrusion made substrate contact across the gap. In controls (not treated with blebbistatin), crossing occurred readily for contacts on LN and after a delay for contacts on FN, but was not observed for contacts on PLO. Thus, f-actin and dynamic MTs are necessary for axon elongation (Chia et al., 2016), but MII inactivation is required for advance from LN or FN onto PLO. Growth cone crossing of a non-adhesive gap revealed the initial steps of directed growth cone advance: 1. Protrusive activity at the leading edge (driven by actin polymerization), 2. adhesive contact with a substrate that supports adhesion-cytoskeletal coupling (i.e, weakening of actomyosin restraint of microtubule advance into a protrusion), 3. microtubule penetration of a protrusion, 4. stabilization of the protrusion as a consequence of microtubule interaction with a putative adhesion complex, and, then 5. further protrusive activity distal to the adhesion complex. Importantly, one of the initial steps is not traction force pulling as demonstrated by the ability of growth cones to cross and advance on all substrates in the absence of MII activity.

Based upon these results and our previous findings (Turney et al., 2016), we propose that the probability of microtubule entry varies with the degree of restraint associated with each adhering protrusion. In an advancing growth cone, which continually produces new protrusions, the selection of a protrusion for stabilization provides a highly sensitive steering mechanism because protrusions compete for microtubule penetration to determine the direction of advance. In the absence of MII activity, steering is lost because restraint is eliminated and the probability of microtubule entry is unregulated.

The stabilization of a protrusion by invading microtubules is the most likely trigger for directed growth cone advance. The mechanism of stabilization and how stabilization is linked to further actin-dependent protrusive activity distally remain unclear (Chia et al., 2016; Zhou and Cohan, 2004). One possibility is that microtubules deliver components necessary for stabilization of adhesion complexes (Kaverina et al., 2002). Another possibility is that microtubule entry promotes advance of smooth endoplasmic reticulum (SER) and that interaction with the adhesion complex stabilizes both the dynamic microtubules and the SER (Dailey and Bridgman, 1989; Pavez et al., 2019; Zhang and Forscher, 2009). These possibilities are not mutually exclusive. Further work will be required to determine the mechanism of stabilization.

It has recently been shown that local coupling of retrograde flow to growth cone point contact adhesions in Xenopus spinal neurons correlates with the rate of advance (Nichol et al., 2016). This is consistent with our previous finding that vinculin dependent adhesion-cytoskeletal coupling affects retrograde flow rates and is necessary for stimulation of DRG neuron outgrowth by NGF (Turney et al., 2016). Furthermore, it has recently been shown that Rho A regulates axon extension mainly through its effects on MII activity (Dupraz et al., 2019). Together these finding support a critical role for MII- and retrograde flow-dependent restraint of microtubule advance.

An important but perhaps subtle conclusion of our findings is that an actin-rich protrusion does not initiate its own stabilization. Instead we think that microtubule entry selects a protrusion and that the probability of selection is inversely related to the restraint of microtubule invasion (see also (Turney et al., 2016). The significance of this seemingly small difference (i.e., whether or not the protrusion initiates stabilization) is that if microtubules initiate stabilization, it allows us to better explain signal integration. It differs from previous interpretations of the role of microtubules in growth cone turning in that we propose that microtubule dependent stabilization of protrusion is an early step in the steering mechanism that will define the new direction rather than a later event that is only important for consolidation. According to our model (Figure 13), protrusions are extended more or less randomly each with its own level of actin-based restraint (determined by interactions with the local environment). One of these protrusions becomes stabilized by invading microtubules with the probability of selection depending on the level of restraint. Growth cone advance then quickly follows in the direction of the selected protrusion. For this model to be correct, axon elongation would consist of a sequence of steps. Each step would be triggered by selection of a protrusion.

**Figure 13.**
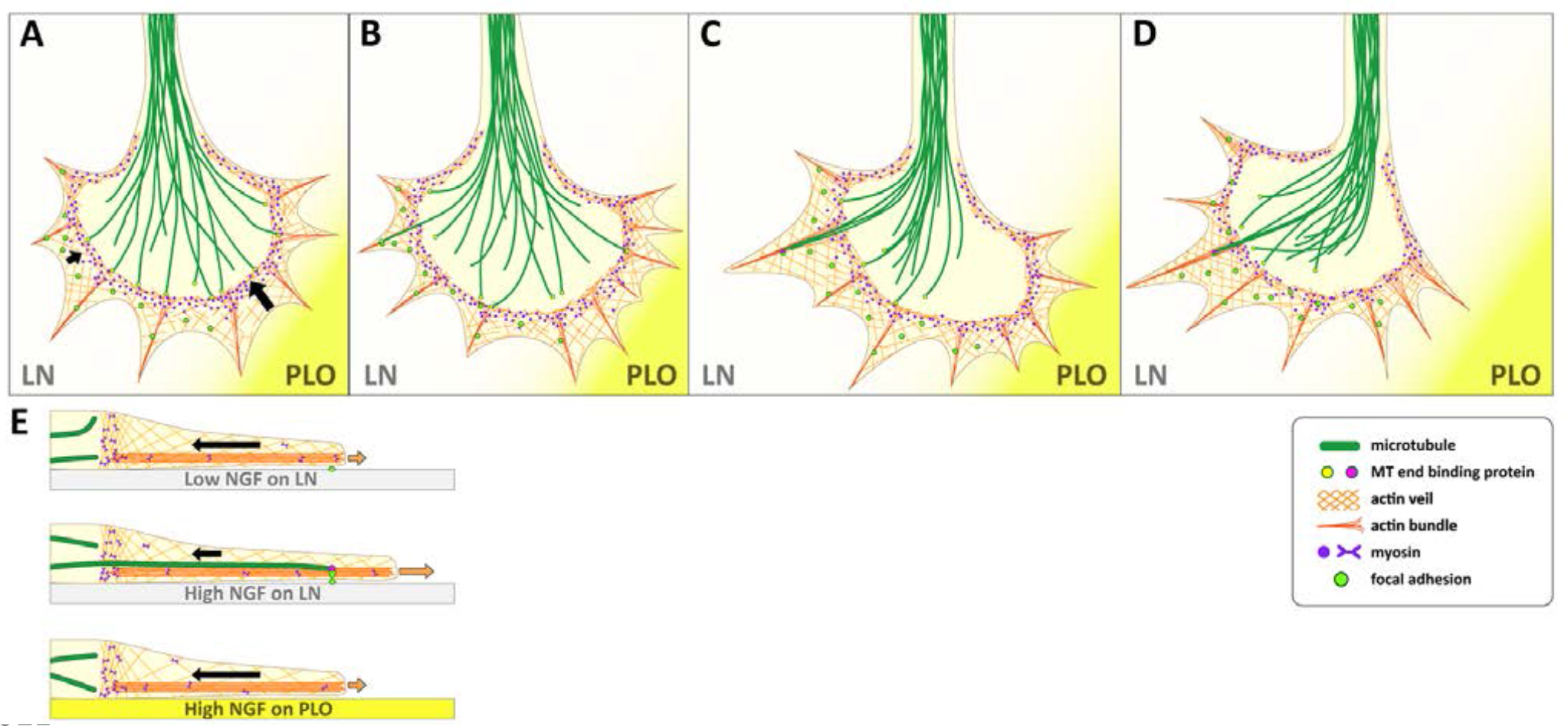
A model of the restraint mechanism of axonal growth cone turning and advance. (A) A growth cone advancing on LN reaches a border with PLO where it extends filopodial protrusions that contact the two different substrates. A transverse arc of actin filaments (cross-linked by MII; arcs are not obvious in rapidly advancing growth cones on LN, but are observed during stopping or pausing) restrains entry of dynamic microtubules (green filaments/yellow end binding proteins) into radial protrusions. Adhesion complexes (green spots) form on LN but not on PLO. As a result, MII-driven retrograde actin flow (black arrows) slows on LN because of adhesion-cytoskeletal coupling on LN but remains fast on PLO. (B) The growth cone continues to extend and retract protrusions more or less randomly while it is stopped at the border. Dynamic microtubules are splayed within the growth cone. The transverse actin bundles and retrograde actin flow actively block microtubule advance into the periphery. One microtubule succeeds (probabilistically) in invading a protrusion on LN due to the slower retrograde actin flow on LN. The actin bundle within the protrusion guides its advance. C) Interaction with the adhesion complex stabilizes the microtubule (represented by the change in end-binding proteins) and the protrusion. Additional microtubules follow. Actin polymerization-driven protrusion is stimulated beyond the adhesion complex. (D) Microtubules progressively assemble into polarized array that becomes bundled in the direction of the stabilized protrusion. Actin protrusion increases beyond the adhesion site. (E) Side view of a filopodium on LN in low NGF: weak adhesion-cytoskeletal coupling results in fast retrograde flow (large black arrow) that sweeps back dynamic microtubules. Side view of a filopodium on LN in high NGF: strong adhesion-cytoskeletal coupling leads to slowing of retrograde flow (small black arrow). Dynamic microtubules more readily penetrate up to the adhesion site. Actin polymerization driven protrusion proceeds beyond adhesion site. Side view of a filopodium on PLO in high NGF: lack of adhesion-cytoskeletal coupling results in a high rate of retrograde flow (large black arrow). Microtubule advance is impeded.

Axon guidance can be understood as a process in which growth cones: 1) integrate signals from many different factors, 2) transduces the signals into a behavior to stay put or advance in a certain direction. A question is whether this process updates continuously or in discrete steps. The latter would suggest that axon locomotive behavior is intrinsically stepwise in nature. This possibility is supported by studies showing that elongation can be modeled as a biased random walk (Katz et al., 1984). In time-lapse recordings axons are seen to elongate in straight runs interrupted by occasional pauses and turns. Furthermore, upon closer examination growth cones zigzag slightly during a straight run. The possibility of stepwise locomotion is also consistent with current modeling of the mechanism of growth cone sensing and steering. The most widely accepted model is that an actin-rich protrusion is extended in the direction of future growth and then invading microtubules reinforce growth in the new direction (Lee and Suter, 2008; Lowery and Van Vactor, 2009). Nevertheless, the case for growth cone movement being stepwise is far from conclusive as can be seen from the fact that theoretical modeling of pathfinding behavior is presented without reference to a motility mechanism (Goodhill and Urbach, 1999; Kobayashi et al., 2010). One reason is that growth cone advance in fast growing axons appears to be continuous not saltatory. Another reason is that the current modeling of growth cone sensing and steering does not reinforce the idea of stepwise advance partly because, according to the proposed mechanism, selection is initiated by the formation of a protrusion. A growth cone often extends many short-lived protrusions before it is seen to turn and/or advance. Thus, it has been unclear how protrusions sense the environment and compete with each other such that only one becomes stabilized. For example slowly advancing or paused growth cones at decision points often have large spread areas and appear to be integrating signals and undergoing cytoskeletal reorganization that gradually lead to advance in a new direction (or a continuation in the same direction) (Mason and Erskine, 2000; Mason and Wang, 1997). Indeed, the well-known turning assay developed by Poo and colleagues analyzes turning as variable tilt of the distal axon/growth cone slightly toward an attractant or away from a repellent (Hopker et al., 1999; Lohof et al., 1992; Zheng et al., 1996).

Our proposed model described above, offers a coherent framework for understanding a number of well-known guidance phenomena. For example, attractant and repellent cues may act by decreasing and increasing restraint, respectively. In other words, attractant cues increase the probability of microtubule invasion of a protrusion, whereas repellent cues decrease the probability. Similarly, neurotrophins may induce faster elongation by causing decreased restraint and consequently a higher rate of protrusion selection. Thus, growth cone turning and advance could be characterized as two aspects of the same growth cone behavior. Finally signal integration can be understood in terms of how different factors exert their effects on restraint. Some factors may influence restraint within a single protrusion (locally) and others on the whole growth cone (globally). The selection process may also be biased by factors that regulate actin or microtubule polymerization (Buck and Zheng, 2002); however, we think that growth cone navigation is dominated by the restraint mechanisms because in the absence of MII activity, elongation continues, but is slower and undirected (Turney and Bridgman, 2005).

The significance of the restraint-based mechanism is perhaps easier to understand if the model is expressed using concepts from evolutionary dynamics (Lewontin, 1974; Taylor et al., 2004). In this model protrusive activity is largely random. Protrusions are born (extend) and die (retract) at rates that result in a relatively small population size. Initially protrusions are identical (equally fit). As they encounter different factors in the local environment, each may “mutate” to have a different fitness largely due to changes in restraint. Fitness varies inversely with the level of restraint. The probability of microtubules invading and stabilizing a protrusion (selection) increases with fitness. Thus, the protrusions with highest fitness are the ones most likely to survive. Selection of a protrusion leads to growth cone advance and further protrusive activity (i.e., reproduction). If every protrusion has the same fitness (e.g., a growth cone advancing on a uniform substrate), then growth cone advance will proceed as a constrained random walk along the axis defined by the orientation of bundled microtubules in the axon because protrusions at the leading edge are aligned with the bundled microtubules and therefore more likely to be invaded and stabilized. In other words, a protrusion is selected with a probability proportional to its position on the growth cone multiplied by its fitness. If protrusions on one side of a growth cone have higher fitness than those in front or on the other side (for instance in response to a change in substrate or a guidance cue gradient), then they may be selected more frequently causing a change in the direction of growth. This is a testable model.

In summary, we show that stopping and stepping behaviors are characteristic of elongating axons. We found that the growth substrate (LN) influences these all-or-none behaviors through its effects on MII-dependent restraint of microtubule entry into actin-rich growth cone protrusions. We observed that stepwise growth cone advance occurs only when microtubules stably invade an actin-rich protrusion. MII-dependent retrograde flow is linked to the probability of microtubule entry. Aversive turning or stopping occurs when the probability of microtubule entry is comparatively low for protrusions extended in the direction of axon growth. From these results we present a model that provides a framework for understanding how mammalian axons navigate through complex environments in vivo. This modeling has immediate implications for discovering conditions under which axon growth will be facilitated or blocked such as during axon regeneration. Furthermore, we think the stochastic mechanism helps explain the robustness of axon pathfinding and suggests that maps are not necessarily constructed from highly specific sets of cues that are in rigidly defined patterns. Rather the maps are likely to be flexible such that perturbation in cues does not render them useless.

## Materials and Methods

### Cell culture and microfluidics device

DRG or SCG neurons from E13.5-14.5 mouse embryos were cultured in microfluidic Campenot chambers using methods described previously (Turney et al., 2016) (Bridgman et al., 2001). The microfluidic chambers were made of polydimethylsiloxane (PDMS) bonded to glass coverslips. Cells were plated on the coated coverslips (0.1 mg/ml PLO+ 20 µg/ml LN or 50 µg/ml FN) following dissociation in the open central wells (2 mm diameter) of microfluidic Campenot chambers. Each microfluidics device contained four Campenot chambers and up to two devices could be bonded to a single coverslip (35 cm in diameter). Each well connected to an open axon chamber by 50, 10 µm wide channels (∼1.0 mm in length). Axon chamber borders between coatings were created using PDMS masks as previously described (Turney and Bridgman, 2005). Unless indicated, drugs were added only to the axon chamber (exchanged as necessary at 24 h intervals). Typically to avoid slow growth through channels, drugs were added only after growth cones entered the axon chamber, but prior to reaching a lane gap. Campenot chambers rely upon fluid pressure to prevent solutes in the axon compartment from reaching the cell body compartment. In an open well system, this is achieved by keeping the fluid level higher in the cell body compartment than in the axon compartment. The small fluid volumes associated with the Campenot chambers necessitated designing a chamber to allow unrestricted high-resolution views of the cells for long times (up to 38 h) on the microscope while preventing evaporation. Our new chamber design is compatible with a stage mounted environmental chamber with heating and CO2. The design is covered by a patent (US Patent 9,939,424) and will be described in detail elsewhere. Each experimental condition was run in duplicate along with two matching controls. Experiments were repeated three times.

### Substrate patterning

The glass surface in the axon compartments of the Campenot chamber systems with the different coatings (LN, FN, PLO) were patterned by region of interest (ROI) scanning using a mode-locked multiphoton laser (800 nm light) on an inverted multiphoton microscope (Zeiss LSM 510 NLO). The lanes created by patterning were aligned with the channels of the microfluidic Campenot chambers and were 9-12 µm in width. Lane patterns were usually created prior to plating cells, but their length could be extended as needed as axons grew in length. Gaps within the lanes were 8-24 µm in width. Gaps were usually created after axons entered lanes since their position was determined by the entry of axons into a lane and how far they extended along the lane. This allowed capture of interactions with gaps within a reasonable time window. The patterning method (referred to as Live Cell Substrate Patterning or LCSP; US patent 8,921,283) will be described in detail elsewhere.

### Transfection

Dissociated cells were transfected using electroporation (Amaxa Nucleofector) prior to plating in the cell compartment of the Campenot chamber. GFP-EB3 was a gift of Niels Galjart; GFP-LifeAct and Ruby-LifeAct were gifts of Dorothy Schafer.

### Antibody/Fluorescence Labeling

Dynamic microtubules in fixed preparations were detected using a Mab to tyrosinated tubulin (Rochlin et al., 1996). Actin in fixed preparations was detected by staining with Rhodamine phalloidin (Sigma).

### Imaging

Growing axons were imaged on a Zeiss LSM 510 NLO using our custom culture chamber in a stage-mounted environmental chamber (PCO) or on an inverted Olympus IX70 equipped with a sensitive CCD camera (Sensicam), LED illumination (Prizmatix) and custom environmental chamber. Extended time lapse imaging on the Zeiss LSM 510 was performed using the Multitime Series macro. To capture gap crossing events growth cones were imaged over hours at regular intervals using combined DIC and reflected light optics (at either 800 or 633 nm).

When crossing appeared imminent in growth cones expressing fluorescent proteins, we switched to combined DIC, reflected light and fluorescence (at the appropriate fluorescence wavelength) during the cross or attempted cross. Time-lapse intervals were varied between 5 sec and 5 min. Ruby-LifeAct was used to monitor actin dynamics following blebbistatin treatment. Reflected light and DIC imaging of blebbistatin treated cultures was always done at 800 nm to avoid phototoxicity (Kolega, 2004). Fixed cultures were imaged either on the Olympus IX70 or the Zeiss LSM 510. Rotary shadowing EM was performed as previously described (Bridgman, 2002). Imaging was done using either a JOEL 1200EX or a JEM-1400.

### Statistical Analysis

Descriptive statistical analysis was carried out using Excel (Microsoft). The ANOVA and unpaired Student’s *t* test was used to determine the significance between two groups.

## ACKNOWLEDGEMENTS

We thank Janet Iwasa for creating the illustration for Figure 8. This work was supported by grants to P.C. Bridgman from NIH (R21 MH081260, R21EB9776) and in part by the Bakewell Neuroimaging Core, supported by the Bakewell Family Foundation and the National Institutes of Health Neuroscience Blueprint Interdisciplinary Center Core Grant P30 (NS057105) to Washington University. The Office of Technology Management at WUSM provided support for patent applications. S. Turney receives support from the Department of Molecular and Cellular Biology and Jeff Lichtman. The fabrication of microfluidics devices was performed in part at the Center for Nanoscale Systems, a member of the National Nanotechnology Infrastructure Network, which is supported by the National Science Foundation under award No. ECS-0335765. The Center for Nanoscale Systems is part of the Faculty of Arts and Sciences at Harvard University. R.M.R. received a postdoctoral fellowship (1 F32 NS60356-01) from the National Institutes of Health.

## SUPPLEMENTAL MOVIES

**Movie S1**. Filopodia dynamics in a growth cone stopped at a non-adhesive block. Images are at 2 min intervals (60 frames) (corresponds to Figure 1C).

**Movie S2**. Growth cone on a LN path crossing a non-adhesive gap during treatment with blebbistatin (corresponds to Figure 2B). Images are at 4 min intervals.

**Movie S3.** Actin dynamics revealed by GFP-LA in an untreated control growth cone on LN (corresponds to Figure 8C, left panel). Peripheral actin-rich protrusion formation with associated retrograde flow occurs mainly on the branch to the left. Images are at 5 sec intervals.

**Movie S4.** Actin dynamics revealed by Ruby-LA in a bleb treated growth cone on LN (corresponds to Figure 8C, right panel). Protrusions are transient. Retrograde flow appears in the peripheral processes, filopodia and proximal neurite. Images are at 5 sec intervals.

**Movie S5**. GFP-EB3 dynamics in a growth cone stopped at a LN-PLO border (corresponds to Figure 10). EB3 comets do not enter into filopodia extending onto PLO, but do enter into protrusions extending onto the non-adhesive border on the right that creates the lane. Images are at 5 sec intervals.

**Movie S6**. GFP-EB3 dynamics in a growth cone on a LN path during crossing of a non-adhesive gap (corresponds to Figure 12). Images are at 5 sec intervals.

## REFERENCES

1. Allingham, J.S., Smith, R., and Rayment, I. (2005). The structural basis of blebbistatin inhibition and specificity for myosin II. Nat Struct Mol Biol 12, 378–379.

2. Athamneh, A.I.M., He, Y., Lamoureux, P., Fix, L., Suter, D.M., and Miller, K.E. (2017). Neurite elongation is highly correlated with bulk forward translocation of microtubules. Scientific reports 7, 7292.

3. Bentley, D., and Toroian-Raymond, A. (1986). Disoriented pathfinding by pioneer neurone growth cones deprived of filopodia by cytochalasin treatment. Nature 323, 712–715.

4. Bridgman, P.C. (2002). Growth cones contain myosin II bipolar filament arrays. Cell Motil Cytoskeleton 52, 91–96.

5. Bridgman, P.C., Dave, S., Asnes, C.F., Tullio, A.N., and Adelstein, R.S. (2001). Myosin IIB is required for growth cone motility. J Neurosci 21, 6159–6169.

6. Buck, K.B., and Zheng, J.Q. (2002). Growth cone turning induced by direct local modification of microtubule dynamics. J Neurosci 22, 9358–9367.

7. Burnette, D.T., Manley, S., Sengupta, P., Sougrat, R., Davidson, M.W., Kachar, B., and Lippincott-Schwartz, J. (2011). A role for actin arcs in the leading-edge advance of migrating cells. Nat Cell Biol 13, 371–381.

8. Burnette, D.T., Schaefer, A.W., Ji, L., Danuser, G., and Forscher, P. (2007). Filopodial actin bundles are not necessary for microtubule advance into the peripheral domain of Aplysia neuronal growth cones. Nat Cell Biol 9, 1360–1369.

9. Challacombe, J.F., Snow, D.M., and Letourneau, P.C. (1997). Dynamic microtubule ends are required for growth cone turning to avoid an inhibitory guidance cue. J Neurosci 17, 3085–3095.

10. Chia, J.X., Efimova, N., and Svitkina, T.M. (2016). Neurite outgrowth is driven by actin polymerization even in the presence of actin polymerization inhibitors. Mol Biol Cell.

11. Dailey, M.E., and Bridgman, P.C. (1989). Dynamics of the endoplasmic reticulum and other membranous organelles in growth cones of cultured neurons. J Neurosci 9, 1897–1909.

12. Davenport, R.W., Dou, P., Rehder, V., and Kater, S.B. (1993). A sensory role for neuronal growth cone filopodia. Nature 361, 721–724.

13. Dent, E.W., Kwiatkowski, A.V., Mebane, L.M., Philippar, U., Barzik, M., Rubinson, D.A., Gupton, S., Van Veen, J.E., Furman, C., Zhang, J., et al. (2007). Filopodia are required for cortical neurite initiation. Nat Cell Biol 9, 1347–1359.

14. Dupraz, S., Hilton, B.J., Husch, A., Santos, T.E., Coles, C.H., Stern, S., Brakebusch, C., and Bradke, F. (2019). RhoA Controls Axon Extension Independent of Specification in the Developing Brain. Curr Biol.

15. Fischer, M., Haase, I., Wiesner, S., and Muller-Taubenberger, A. (2006). Visualizing cytoskeleton dynamics in mammalian cells using a humanized variant of monomeric red fluorescent protein. FEBS Lett 580, 2495–2502.

16. Goeckeler, Z.M., Bridgman, P.C., and Wysolmerski, R.B. (2008). Nonmuscle myosin II is responsible for maintaining endothelial cell basal tone and stress fiber integrity. Am J Physiol Cell Physiol 295, C994–C1006.

17. Gomez, T.M., Roche, F.K., and Letourneau, P.C. (1996). Chick sensory neuronal growth cones distinguish fibronectin from laminin by making substratum contacts that resemble focal contacts. J Neurobiol 29, 18–34.

18. Gomez, T.M., and Spitzer, N.C. (1999). In vivo regulation of axon extension and pathfinding by growth-cone calcium transients. Nature 397, 350–355.

19. Goodhill, G.J., and Urbach, J.S. (1999). Theoretical analysis of gradient detection by growth cones. J Neurobiol 41, 230–241.

20. Hopker, V.H., Shewan, D., Tessier-Lavigne, M., Poo, M., and Holt, C. (1999). Growth-cone attraction to netrin-1 is converted to repulsion by laminin-1. Nature 401, 69–73.

21. Hur, E.M., Yang, I.H., Kim, D.H., Byun, J., Saijilafu, Xu, W.L., Nicovich, P.R., Cheong, R., Levchenko, A., Thakor, N., et al. (2011). Engineering neuronal growth cones to promote axon regeneration over inhibitory molecules. Proc Natl Acad Sci U S A 108, 5057–5062.

22. Kahn, O.I., and Baas, P.W. (2016). Microtubules and Growth Cones: Motors Drive the Turn. Trends Neurosci 39, 433–440.

23. Kater, S.B., and Rehder, V. (1995). The sensory-motor role of growth cone filopodia. Curr Opin Neurobiol 5, 68–74.

24. Katz, M.J., George, E.B., and Gilbert, L.J. (1984). Axonal elongation as a stochastic walk. Cell Motil 4, 351–370.

25. Kaverina, I., Krylyshkina, O., and Small, J.V. (2002). Regulation of substrate adhesion dynamics during cell motility. Int J Biochem Cell Biol 34, 746–761.

26. Ketschek, A.R., Jones, S.L., and Gallo, G. (2007). Axon extension in the fast and slow lanes: substratum-dependent engagement of myosin II functions. Dev Neurobiol 67, 1305–1320.

27. Kobayashi, T., Terajima, K., Nozumi, M., Igarashi, M., and Akazawa, K. (2010). A stochastic model of neuronal growth cone guidance regulated by multiple sensors. J Theor Biol 266, 712–722.

28. Kolega, J. (2004). Phototoxicity and photoinactivation of blebbistatin in UV and visible light. Biochem Biophys Res Commun 320, 1020–1025.

29. Lamoureux, P., Buxbaum, R.E., and Heidemann, S.R. (1989). Direct evidence that growth cones pull. Nature 340, 159–162.

30. Lee, A.C., and Suter, D.M. (2008). Quantitative analysis of microtubule dynamics during adhesion-mediated growth cone guidance. Dev Neurobiol 68, 1363–1377.

31. Lewontin, R.C. (1974). The Genetic Basis of Evolutionary Change (New York: Columbia University Press).

32. Limouze, J., Straight, A.F., Mitchison, T., and Sellers, J.R. (2004). Specificity of blebbistatin, an inhibitor of myosin II. J Muscle Res Cell Motil 25, 337–341.

33. Lohof, A.M., Quillan, M., Dan, Y., and Poo, M.M. (1992). Asymmetric modulation of cytosolic cAMP activity induces growth cone turning. J Neurosci 12, 1253–1261.

34. Loudon, R.P., Silver, L.D., Yee, H.F., Jr., and Gallo, G. (2006). RhoA-kinase and myosin II are required for the maintenance of growth cone polarity and guidance by nerve growth factor. J Neurobiol 66, 847–867.

35. Lowery, L.A., and Van Vactor, D. (2009). The trip of the tip: understanding the growth cone machinery. Nat Rev Mol Cell Biol 10, 332–343.

36. Mason, C., and Erskine, L. (2000). Growth cone form, behavior, and interactions in vivo: retinal axon pathfinding as a model. J Neurobiol 44, 260–270.

37. Mason, C.A., and Wang, L.C. (1997). Growth cone form is behavior-specific and, consequently, position-specific along the retinal axon pathway. J Neurosci 17, 1086–1100.

38. Medeiros, N.A., Burnette, D.T., and Forscher, P. (2006). Myosin II functions in actin-bundle turnover in neuronal growth cones. Nat Cell Biol 8, 215–226.

39. Menon, S., Boyer, N.P., Winkle, C.C., McClain, L.M., Hanlin, C.C., Pandey, D., Rothenfusser, S., Taylor, A.M., and Gupton, S.L. (2015). The E3 Ubiquitin Ligase TRIM9 Is a Filopodia Off Switch Required for Netrin-Dependent Axon Guidance. Dev Cell 35, 698–712.

40. Mitchison, T., and Kirschner, M. (1988). Cytoskeletal dynamics and nerve growth. Neuron 1, 761–772.

41. Mortimer, D., Fothergill, T., Pujic, Z., Richards, L.J., and Goodhill, G.J. (2008). Growth cone chemotaxis. Trends Neurosci 31, 90–98.

42. Nadar, V.C., Ketschek, A., Myers, K.A., Gallo, G., and Baas, P.W. (2008). Kinesin-5 is essential for growth-cone turning. Curr Biol 18, 1972–1977.

43. Nichol, R.H.t., Hagen, K.M., Lumbard, D.C., Dent, E.W., and Gomez, T.M. (2016). Guidance of Axons by Local Coupling of Retrograde Flow to Point Contact Adhesions. J Neurosci 36, 2267–2282.

44. Pavez, M., Thompson, A.C., Arnott, H.J., Mitchell, C.B., D’Atri, I., Don, E.K., Chilton, J.K., Scott, E.K., Lin, J.Y., Young, K.M., et al. (2019). STIM1 Is Required for Remodeling of the Endoplasmic Reticulum and Microtubule Cytoskeleton in Steering Growth Cones. J Neurosci 39, 5095–5114.

45. Riedl, J., Crevenna, A.H., Kessenbrock, K., Yu, J.H., Neukirchen, D., Bista, M., Bradke, F., Jenne, D., Holak, T.A., Werb, Z., et al. (2008). Lifeact: a versatile marker to visualize F-actin. Nat Methods 5, 605–607.

46. Robles, E., and Gomez, T.M. (2006). Focal adhesion kinase signaling at sites of integrin-mediated adhesion controls axon pathfinding. Nat Neurosci 9, 1274–1283.

47. Rochlin, M.W., Dailey, M.E., and Bridgman, P.C. (1999). Polymerizing microtubules activate site-directed F-actin assembly in nerve growth cones. Mol Biol Cell 10, 2309–2327.

48. Rochlin, M.W., Wickline, K.M., and Bridgman, P.C. (1996). Microtubule stability decreases axon elongation but not axoplasm production. J Neurosci 16, 3236–3246.

49. Sabry, J.H., O’Connor, T.P., Evans, L., Toroian-Raymond, A., Kirschner, M., and Bentley, D. (1991). Microtubule behavior during guidance of pioneer neuron growth cones in situ. J Cell Biol 115, 381–395.

50. Schaefer, A.W., Kabir, N., and Forscher, P. (2002). Filopodia and actin arcs guide the assembly and transport of two populations of microtubules with unique dynamic parameters in neuronal growth cones. J Cell Biol 158, 139–152.

51. Schaefer, A.W., Schoonderwoert, V.T., Ji, L., Mederios, N., Danuser, G., and Forscher, P. (2008). Coordination of actin filament and microtubule dynamics during neurite outgrowth. Dev Cell 15, 146–162.

52. Stepanova, T., Slemmer, J., Hoogenraad, C.C., Lansbergen, G., Dortland, B., De Zeeuw, C.I., Grosveld, F., van Cappellen, G., Akhmanova, A., and Galjart, N. (2003). Visualization of microtubule growth in cultured neurons via the use of EB3-GFP (end-binding protein 3-green fluorescent protein). J Neurosci 23, 2655–2664.

53. Stephens, L., Milne, L., and Hawkins, P. (2008). Moving towards a better understanding of chemotaxis. Curr Biol 18, R485–494.

54. Straight, A.F., Cheung, A., Limouze, J., Chen, I., Westwood, N.J., Sellers, J.R., and Mitchison, T.J. (2003). Dissecting temporal and spatial control of cytokinesis with a myosin II Inhibitor. Science 299, 1743–1747.

55. Suter, D.M., and Forscher, P. (2001). Transmission of growth cone traction force through apCAM-cytoskeletal linkages is regulated by Src family tyrosine kinase activity. J Cell Biol 155, 427–438.

56. Suter, D.M., and Miller, K.E. (2011). The emerging role of forces in axonal elongation. Progress in neurobiology 94, 91–101.

57. Tanaka, E.M., and Kirschner, M.W. (1991). Microtubule behavior in the growth cones of living neurons during axon elongation. J Cell Biol 115, 345–363.

58. Taylor, C., Fudenberg, D., Sasaki, A., and Nowak, M.A. (2004). Evolutionary game dynamics in finite populations. Bull Math Biol 66, 1621–1644.

59. Turney, S.G., Ahmed, M., Chandrasekar, I., Wysolmerski, R.B., Goeckeler, Z.M., Rioux, R.M., Whitesides, G.M., and Bridgman, P.C. (2016). Nerve growth factor stimulates axon outgrowth through negative regulation of growth cone actomyosin restraint of microtubule advance. Mol Biol Cell 27, 500–517.

60. Turney, S.G., and Bridgman, P.C. (2005). Laminin stimulates and guides axonal outgrowth via growth cone myosin II activity. Nat Neurosci 8, 717–719.

61. Van Goor, D., Hyland, C., Schaefer, A.W., and Forscher, P. (2012). The role of actin turnover in retrograde actin network flow in neuronal growth cones. PLoS One 7, e30959.

62. Waterman-Storer, C.M., Worthylake, R.A., Liu, B.P., Burridge, K., and Salmon, E.D. (1999). Microtubule growth activates Rac1 to promote lamellipodial protrusion in fibroblasts. Nat Cell Biol 1, 45–50.

63. Witte, H., Neukirchen, D., and Bradke, F. (2008). Microtubule stabilization specifies initial neuronal polarization. J Cell Biol 180, 619–632.

64. Yang, Q., Zhang, X.F., Pollard, T.D., and Forscher, P. (2012). Arp2/3 complex-dependent actin networks constrain myosin II function in driving retrograde actin flow. J Cell Biol 197, 939–956.

65. Yvon, A.M., Wadsworth, P., and Jordan, M.A. (1999). Taxol suppresses dynamics of individual microtubules in living human tumor cells. Mol Biol Cell 10, 947–959.

66. Zhang, X.F., and Forscher, P. (2009). Rac1 modulates stimulus-evoked Ca(2+) release in neuronal growth cones via parallel effects on microtubule/endoplasmic reticulum dynamics and reactive oxygen species production. Mol Biol Cell 20, 3700–3712.

67. Zheng, J.Q., Wan, J.J., and Poo, M.M. (1996). Essential role of filopodia in chemotropic turning of nerve growth cone induced by a glutamate gradient. J Neurosci 16, 1140–1149.

68. Zhou, F.Q., and Cohan, C.S. (2004). How actin filaments and microtubules steer growth cones to their targets. J Neurobiol 58, 84–91.

